# “A Proteogenomic workflow reveals distinct molecular phenotypes related to breast cancer appearance”

**DOI:** 10.1101/2020.05.05.077974

**Authors:** Tommaso De Marchi, Paul Theodor Pyl, Martin Sjöstrom, Stina Klasson, Hanna Sartor, Johan Malmström, Lars Malmström, Emma Niméus

**Author notes:** These authors equally contributed to this study (Tommaso De Marchi, Paul Theodor Pyl). **Corresponding authors** Tommaso De Marchi, Emma Niméus.

## Abstract

Proteogenomics approaches have enabled the generation of extensive information levels when compared to single omics technology studies, although burdened by massive experimental efforts. Here, we developed four improvements of a data independent acquisition mass spectrometry proteogenomics workflow to reveal distinct molecular phenotypes related to breast cancer appearance. We confirm mutational processes detectable at the protein level and highlight quantitation and pathway complementarity between RNA and protein data. Our analyses also validated previously established enrichments of estrogen receptor-dependent molecular features relating to transcription factor expression, and provided evidence for molecular differences related to the presence of mammographic appearances in spiculated tumors. In addition, several transcript-protein pairs displayed radically different abundance correlations depending on the overall clinical and pathological properties of the tumor. These results demonstrate that there are differentially regulated protein networks in clinically relevant sample groups, and that these protein networks influence both cancer biology as well as the abundance of potential biomarkers and drug targets.

## Introduction

Breast cancer (BC) is the most common female malignancy that affects women. Despite the constant increase in incidence rate, BC mortality is steadily decreasing due to better care, the availability of new treatment options, and a deeper understanding of the mutational and molecular dynamics of each breast cancer type (DeSantis *et al*, 2019; Fachal *et al*, 2020). BCs are generally classified according to the status of four markers: Estrogen and Progesterone Receptors (ER and PgR), the receptor Tyrosine kinase ERBB2, and Ki67. Subsequent advances explored the molecular heterogeneity of BCs by mRNA analysis defining several intrinsic subtypes (Perou *et al*, 2000), which in turn helped in defining patient prognosis and treatment strategies (Coates *et al*, 2015). In addition, appearance and histological features of breast tumors such as grade of differentiation have been shown to associate to receptor status and specific molecular subtypes (Schnitt, 2010).

Mammographic imaging plays an important role in breast cancer detection and is especially relevant for early tumor detection. Mammographic techniques and improved image analysis have revealed that a breast cancer can exhibit different mammographic appearances, such as spiculations. Mammographic spiculated tumors possess a star-like appearance, which is an indicator of invasiveness linked to cancer infiltration and fibrotic growth around the tumor (Sartor *et al*, 2015; Alexander *et al*, 2006). On the molecular level, spiculated tumors have been shown to be enriched in ER and PgR positive tumor group, often comprised within the Luminal A molecular subtype, and have been linked to better prognosis when compared to well-defined and micro-calcified masses (Evans *et al*, 2006; Taneja *et al*, 2008; Bullier *et al*, 2013). These findings indicate that there are both receptor status (and intrinsic subtypes) dependent and independent molecular drivers that contribute to the spiculated appearance, although the relationship between mutational events and downstream protein regulations responsible for spiculation has remained uncharacterized.

Previous breast cancer reports have demonstrated that proteogenomics, the integration of genomic/transcriptomic and proteomic data, provides a high degree of complementarity for improved definition of molecular drivers in breast cancer. For example, these studies have identified protein-level evidence of genomic aberrations such as chromosomal losses, defined new regulation clusters such as G-protein coupled receptors, identified potential new antigens for immunotherapy, and investigated the discrepancies between transcript and protein pairs in the context of molecular pathways (e.g. metabolism, coagulation) (Mertins *et al*, 2016; Johansson *et al*, 2019; Tyanova *et al*, 2016a). Altogether, these studies revealed that the integration of genomic/transcriptomic and proteomic features not only achieves a degree of complementarity for a better definition of cancer biological characteristic, but also indicate that transcript and protein discrepancies are related to tumor molecular subtype and protein class. In contrast, tumor context-dependent features and their influence on RNA and protein abundance have been sparsely investigated, with little regard on their impact on biomarker measurement, drug target monitoring, or immunotherapy epitope expression.

So far, most proteogenomics studies in BC have relied on peptide-fractionation followed by extensive data dependent acquisition (DDA) MS analysis, typically associated with high instrument usage. However, a recent study employed data independent acquisition (DIA) MS to define high-quality protein maps of BC subtypes, identifying the protein markers INPP4B, CDK1, and ERBB2 as discriminatory of key BC histopathological features, pinpointing biological pathways tied to tumor phenotypes, and assessing expression similarities between transcript and protein abundance (Bouchal *et al*, 2019). These results show that DIA MS constitute an alternative strategy for the quantitative investigation of protein expression in a proteogenomic context, by generating tissue specific spectral libraries and achieving high identification rates out of single-shot analysis. DIA (or SWATH (Gillet *et al*, 2012; Collins *et al*, 2013)) provides consistent peptide/protein identification rates across samples and achieves quantitative precision similar to targeted MS approaches such as selected reaction monitoring (SRM (Teleman *et al*, 2017; Malmström *et al*, 2015; Gillet *et al*, 2012)). Protein identification and quantification is accomplished via DDA-based spectral libraries that play an important role in the reproducible quantification of the breast cancer proteome. In this context, the continuously increasing coverage of the breast cancer proteome achieved by the research community provides new opportunities to further increase and refine breast cancer specific assay libraries, thus improving DIA-based quantification. In addition, the use of spectral libraries previously generated by DDA MS runs has the potential to allow better FDR control for the identification of proteins and their isoforms or mutation-defined single amino acid variants (SAVs). This due to the fact that the data is searched against a smaller database of previously observed peptides rather than whole-proteome search space, impacting the distribution of the identification false discovery rate (FDR) (Malmström *et al*, 2015).

Here, we developed new workflows to improve DIA MS-based proteogenomic analysis, to enable the identification of molecular pathways related to cancer biology and, specifically, associated with mammographic appearances of breast cancer. The improved DIA MS-based workflow coupled to acquired RNA-seq of primary breast cancer tissues exposed transcriptome- and proteome-wide changes dependent on tumor context, and revealed molecular drivers of relevance for receptor status and appearance in breast cancer. The most notable feature was related to the enrichment of factors involved in Epithelial to Mesenchymal Transition (EMT) in spiculated tumors, in line with the concept that protrusions (i.e. spiculae) form the invading front of the cancer and re-model the surrounding healthy breast tissue through for example metalloproteases. Our results not only establish DIA as a viable technique in the field of proteogenomics, but also opens new venues in the elucidation of the biology underneath breast cancer appearance.

## Results

### Generation of a proteogenomics data set for breast cancer

In this study, we selected 21 well-characterized tumor samples from a larger collection (Sjöström *et al*, 2018) of which mammography images, hormonal receptor status and clinical histopathological features were available (**Fig 1A-C** and **Table S1**). Tumor samples were subjected to RNA extraction, and RNA sequencing (RNA-seq) analysis was performed. In addition, we added two proteomic data layers by extracting proteins using ALLPrep (flowthroughs; FT) and standard tissue homogenization (whole tissue lysate; WTL) from the same specimens, followed by trypsin digestion. Peptide fractions were generated via Strong Anion Exchange (SAX) separation prior to DDA MS analysis (as in (Tyanova *et al*, 2016a)), while no fractionation was performed prior DIA MS runs (**Fig 1D**). At the DDA level FT and WTL runs presented potential complementarity, with only minor discrepancies in the number of MS2 events (FT median triggered: 54,059.5, median identified: 6,340; WTL median triggered: 54,465, median identified: 5,571.5) and overall peptide and protein identifications (**Fig S1A-B**). These differences were likely related to ion suppression or matrix effect factors which in turn, as also shown in other studies (De Marchi *et al*, 2016a), impact data density and identification consistency (**Fig S1C-D**). Of interest, the combined subsets achieved higher proteome coverage (**Fig S1E**). On a quantitative level, more than 95% of proteins and around 60% of pathways showed comparable enrichment levels, while only a minority displaying a significant difference (**Fig S2A-B** and **Table S2-3**). While we believed these discrepancies to be due to tumor content in each tissue, no clear association was detected (**Fig S2C**). These two data sets were combined for downstream DIA library generation.

**Figure 1.**
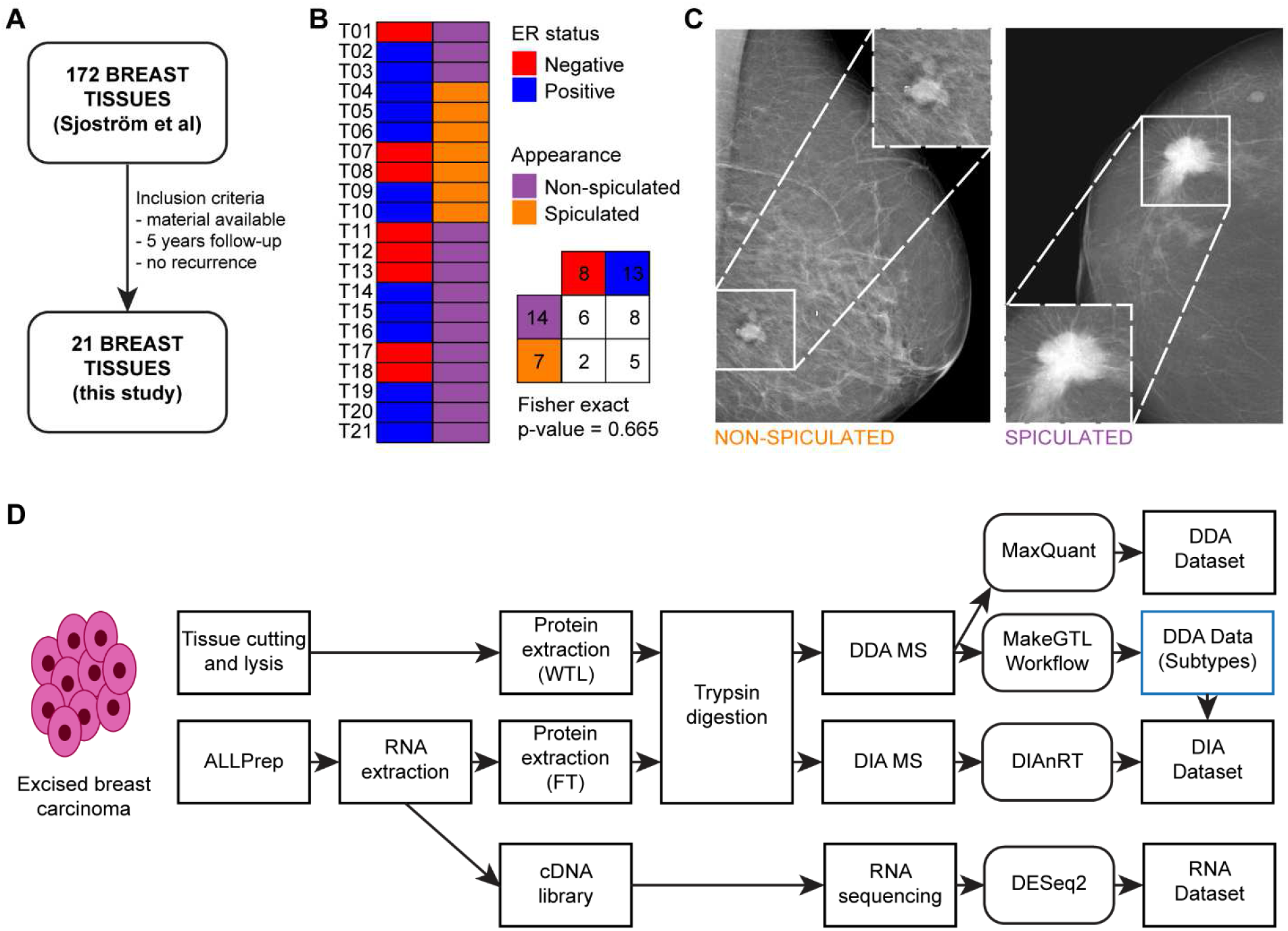
Experimental workflow of this study. A total of 21 samples derived from a larger cohort (N = 172; see Methods) were processed as WTL (MS only) and FT (RNA-seq and MS) for downstream analysis (Panel A). Panel B displays the overlap between the molecular and appearance features evaluated in this study, for which no association as found (Fisher exact p-value = 0.665). Panel C shows examples of non-spiculated and spiculated tumor masses. Panel D displays the experimental workflow of our RNA and MS (DDA and DIA) analyses: tumor tissues were cut into slices and processed by ALLPREP. RNA and protein fractions were extracted and processed from ALLPREP sample preparation for downstream RNA-sequencing and DDA/DIA MS, respectively. Tissue slices were prepared only for downstream MS (DDA/DIA). Samples for DDA were fractionated using Strong Anion Exchange columns (SAX; 6 fractions) so to generate a comprehensive breast cancer library for single-shot DIA MS data searches. Abbreviations: DDA: Data Dependent Acquisition; DIA: Data Independent Acquisition; ER: Estrogen Receptor; FDR: false discovery rate; FT, flowthrough; MS: Mass Spectrometry; RT: Retention Time; WTL: Whole Tissue Lysate.

In the next step, the RNA, DDA, and DIA data layers were used to develop workflows to improve DIA MS-based proteogenomic analysis of breast cancer (**Fig S3A**). We first developed a DIA library creation approach that uses spectral clustering to refine the result and applied it to multiple datasets, i.e. our own DDA data (Ft and WTL combined; De Marchi), as well as DDA data from Tyanova *et al* and Bouchal *et al*. Furthermore, we improved the DIA-MS search workflow by implementing an iterative process for selecting intrinsic RT peptides from the DIA-MS data being searched, thus increasing sensitivity (**Fig S4A-B**). Finally, we integrated transcript sequence information and single nucleotide variants into the sequence used for library creation (**Fig S4C-D**). Using this approach, a transcript-aware DIA library was created, which facilitated the integration of proteomics data with RNA-seq based differential transcript usage information. In parallel to this, a DIA library containing peptides-specific to SAVs (predicted from SNV calling on the RNA-seq data) was created. The combined output from the new workflows resulted in three assay libraries covering 71,152 (this study: De Marchi), 61,282 (Tyanova) and 41,018 (Bouchal) peptide groups (i.e. peptide + charge), which mapped to 9,953, 9,678 and 6,971 proteins, respectively. In addition, the workflows generated 1,025 RNA guided transitions for 74 SAVs and 89,5381 DTU informed transitions covering 43,910 different transcripts matching 12,488 genes. Further, we used the three assay libraries and the more accurate estimation of retention time to improve the peptide identification rates from the DIA experiments. The results from the three libraries were combined to create a superset, in which peptide p-values were conditionally selected for downstream Q-value determination, feature alignment, and re-quantification (**Fig S3A** and **Fig S4A-B**), which resulted in the quantification of 28,746 matching 4,936 proteins (**Fig S3B**). In addition, we analyzed a set of 4 normal breast samples derived from breast reduction surgery, quantifying 7054 peptides mapping 2402 proteins. The low amount of identifications in healthy breast tissue samples likely derives from the low amount of epithelial breast tissue when compared to adipocyte content.

The 28,746 identified peptides out of our search provide information used for protein quantification and to detect presence of SAV and DTU, while the RNA sequencing information provides information of SNV, DTU and RNA abundance. In the final step, the data layers were integrated and compared to extract information of relevance for spiculation and receptor status regarding frequency of DTU, to determine the degree of corroboration or discrepancy between protein and RNA quantitative information, and do identify SAV-specific peptides.

### Comparison between RNA and proteomic data layers

Based on the combined proteogenomics data set, we first compared the overall quantitative measurements between the transcriptomic and proteomic data layers. We matched transcripts to their respective protein products in our proteome datasets to assess the dynamic range of the data layers. The RNA data displayed a flatter dynamic range slope, which might relate to technological differences between gene and protein quantitation technologies, where RNA-seq achieves more even gene quantitation across the transcriptome, or that mRNA data does not accurately reflect post-transcriptional processes, such as ubiquitination and degradation processes that operate exclusively at the protein level (**Fig 2A**). Conversely, the proteomic data layers displayed a wider dynamic range although the transcript and protein abundances are often located in similar quantiles across each sigmoid. In order to more systematically compare transcriptomic and proteomics data layers, we calculated transcript-protein correlations for all detected transcript-protein pairs. We observed a relatively wide range of correlations with a Spearman Rho range of −0.649 to 0.956 (**Fig 2B**), corroborating findings from previous work (Johansson *et al*, 2019). Overall, more than 75% of transcript-protein pairs showed positive correlation coefficients and similar quantitative differences across proteins in the clinically relevant ER status protein group (**Fig 2C**). In contrast, a minority of proteins displayed negative correlations such as RBM39, EXOC3. However, statistical analysis revealed that none of the negative correlations were significant after p-value adjustment, suggesting that the presence of anti-correlating transcript-protein pairs might have a smaller biological effect (**Fig S5A-B**). The high variability of transcript-protein correlations may be dependent on technical limitations or be related to specific protein subclasses and biological pathways, as recently shown in another BC study (Johansson *et al*, 2019). To clarify this, we determined that there was no association observed between transcript-protein pair correlations and signal intensity or sequence coverage (**Fig S6A-D**). In contrast, the degree of correlation was strongly coupled to specific pathways such as RNA splicing or inflammatory response (**Fig 2D-E**). Altogether these results suggest that factors that alter protein abundances specifically, such as post-translational modification and protein degradation have a larger impact on certain protein classes, likely related to cellular regulation of internal processes as well as response to external stimuli, such as mitophagy (Jovanovic *et al*, 2015).

**Figure 2.**
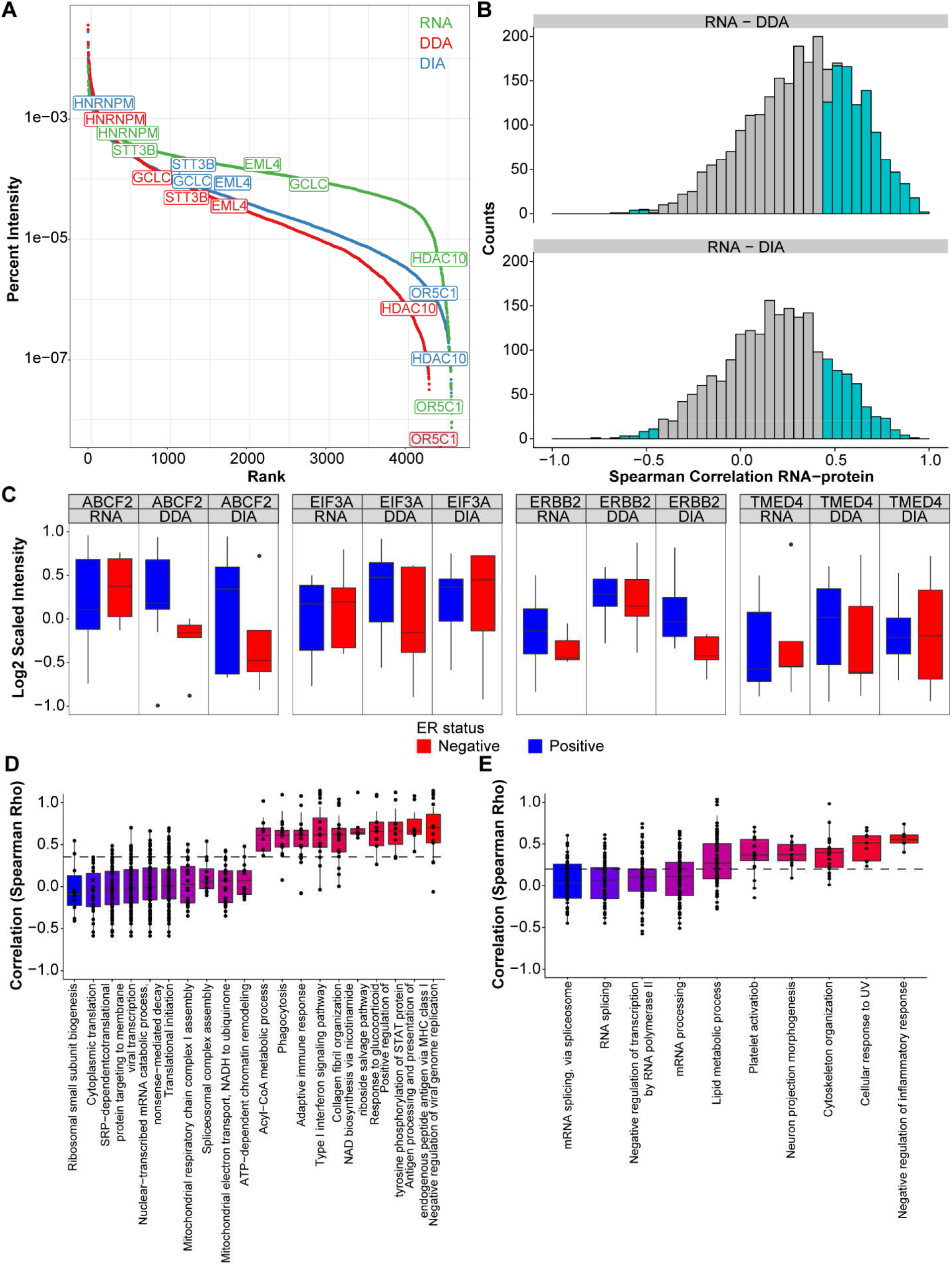
Overall comparison between transcriptomic and proteomic data layers. Panel A displays the dynamic range of transcript and protein intensities of matching identifications in our RNA (green), DDA (blue) and DIA (red) MS data (example of transcript-protein pairs displaying similar abundances across data layers are labeled). Distributions of Spearman correlations between matching transcript and protein (DDA: top; DIA: bottom) abundances are displayed in panel B (gray: non-significant; light blue: significant), while examples of differential expression across ER positive (blue) and negative (red) tumors for positively correlated proteins are shown in panel C. Panel D and E display the distribution of transcript-protein correlations for significant (q-value < 0.15; see Methods for details) GOBP pathways out of our DDA and DIA MS analyses, respectively. Color gradient is representative of low (blue) and high (red) median transcript-protein correlation for each GOBP term. Acronyms: DDA: Data Dependent Acquisition; DIA: Data Independent Acquisition; ER: Estrogen Receptor; GOBP: Gene Ontology Biological Process.

### Pathways related to estrogen receptor status and mammographic appearances

The combined proteogenomics data set further provides new possibilities to investigate differences between clinically relevant tumor groups such as receptor status and appearance features such as spiculation. For this reason, we stratified our breast cancer sample set according to ER status and appearance (**Fig 1B**) and extracted the top 100 most discriminatory genes for each group comparison (**Table S4**-**5**). The direction of regulation was similar for most transcript-protein pairs for both ER status and spiculation (**Fig 3A-B** and **Fig S7A-B** : blue dots), although a few transcript-protein pairs displayed an opposite regulation pattern (**Fig 3A-B** and **Fig S7A-B** : blue dots). In both comparisons, around 70% of transcript-protein pairs displayed similar expression dynamics and ~30% of the pairs showed different regulation direction, although the selected protein-transcript pairs for each comparison was different. However, the magnitude of difference was substantially higher in the ER status group when compared to tumor appearance analysis. This comes as no surprise as ER signaling is one of the strongest signaling pathways in breast cancer. We conclude that although there are differentially regulated transcript-pair produced the spiculated tumors, the magnitude is substantially smaller compared to ER status.

**Figure 3.**
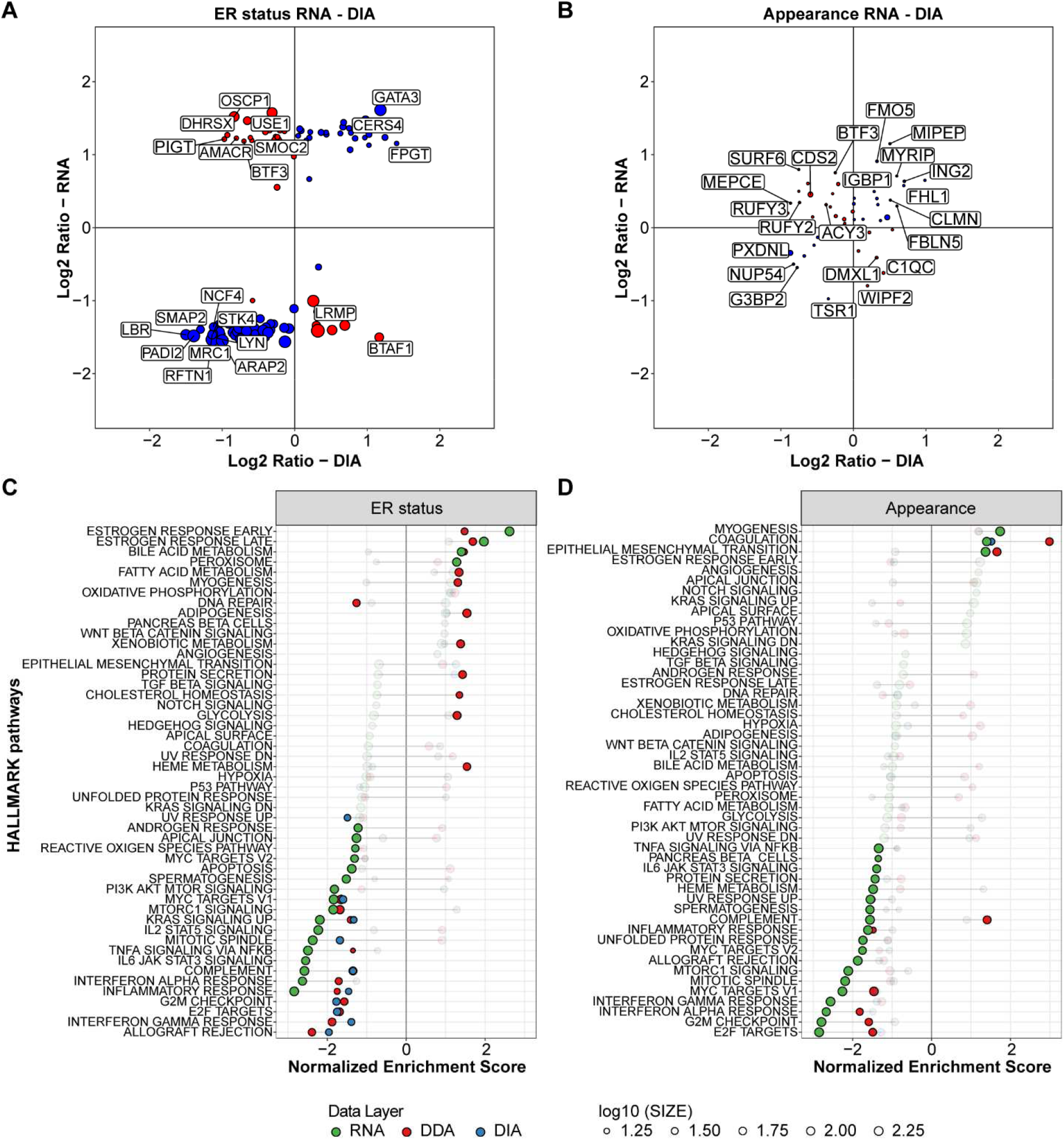
Comparison between transcriptomic and proteomic data in the context of Estrogen Receptor and Appearance statuses. Out of differential expression analyses of RNA-seq data we extracted the most significant transcripts with corresponding detected protein and compared transcript-protein pairs Log2ratios between groups (ER status, panel A; Appearance, panel B). Transcript-protein pairs that displayed similar enrichment at the RNA and protein (DIA) level are marked in blue, while proteins with discrepant enrichment are marked in red. Dot size is relative to adjusted p-value. GSEA analyses were performed on all data layers (RNA, DDA, and DIA) for ER and spiculation statuses, and the top 50 enriched pathways in RNA, DDA, and DIA were compared (Panel C). Left panel displays overlap of GSEA analyses for ER status, while right panel shows results of analysis of appearance features (i.e. spiculation vs no-spiculation). Significant pathways in each data layer (RNA: green; DDA: red; DIA: blue) are marked in full color. Positive scores mark enrichment in ER positive and spiculated tumors, respectively, while negative scores define enrichments in ER negative and non-spiculated samples. Dot size is relative to transcript/protein being found as a core enriched item in GSEA analysis. Acronyms: DDA: Data Dependent Acquisition; DIA: Data Independent Acquisition; ER: Estrogen Receptor; FDR: False Discovery Rate; GSEA: Gene Set Enrichment Analysis.

Next, we investigated whether the combined transcriptomic and proteomic data could provide complementary information of pathways of relevance for the breast cancer subgroups (Mertins *et al*, 2016), by performing GSEA analysis on all three data layers for ER status and appearance (**Fig 3C**). As expected, the estrogen early and late response gene networks were the pathways with highest positive enrichment scores for ER status. For spiculation, the pathways with the highest enrichment score were myogenesis, coagulation and epithelial-mesenchymal transition (EMT). In contrast, immune response (e.g. Allograft Rejection, Interferon Response) and Cell Cycle (e.g. G2M Checkpoint) pathways were found enriched in non-spiculated tumors. Of interest, we note that a higher proportion of pathways out of RNA and protein analyses for ER status was found significant and with the same enrichment directionality. In addition, the proteomic and transcriptomic data layers provide complementarity information regarding pathways related to translation such as E2F targets and immune responses such as interferon gamma response, where significant changes were only detected at the protein level. Collectively, these findings hint at the fact that alterations in specific pathways or class of proteins might only be detected at the protein level.

Of the pathways with the highest enrichment score we selected three for further analysis. The most enriched (by enrichment score and Q-value) pathway in ER positive tumors was the Estrogen Response Early gene set (**Fig 4A**), which includes genes involved in signal transduction processes and cell differentiation (e.g. IGF1R, MUC1), as well as transcription factor-associated proteins such as MED24. Here enriched transcripts and proteins included both ER bound proteins (e.g. FKBP4), and genes activated downstream of ER transcriptional activity (e.g. ABCA3), which relates to downstream activation of breast tissue hormone-dependent proliferation mechanisms.

**Figure 4.**
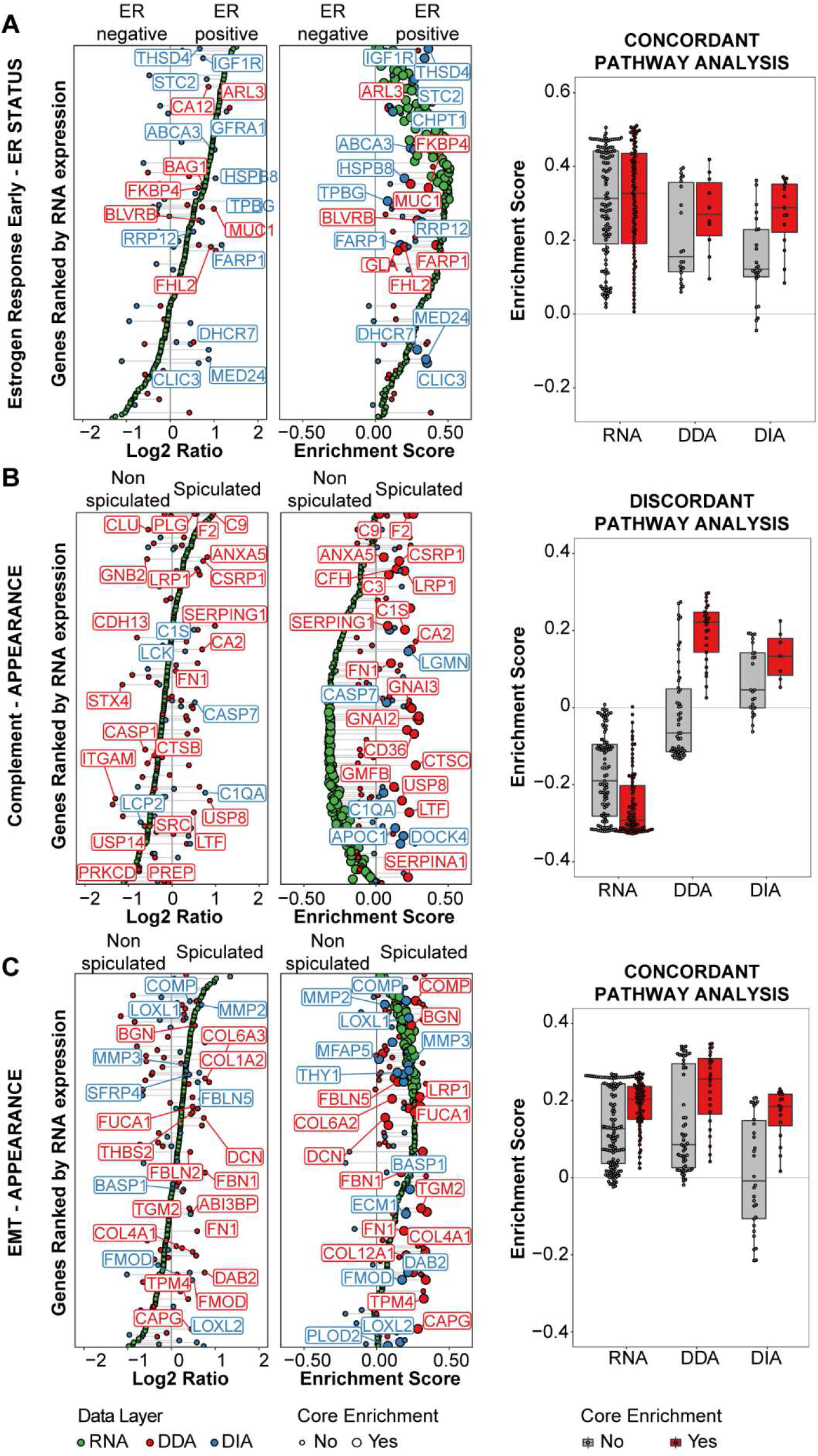
Pathway-level comparison of transcript-protein pairs. Figure displays transcript-protein-wise comparison within significant pathways out of GSEA analyses for ER status (Estrogen Response Early, panel A) and Appearance (Complement, panel B; Epithelial Mesenchymal Transition; panel C). Left panels display Log2Ratios of each transcript/protein (ranked by RNA expression) between ER positive/negative and spiculated/non-spiculated tumors, while center panels display the corresponding enrichment scores in each data layer (RNA: green; DDA: red; DIA: blue). Right panels show distribution of enrichment scores for core enriched (red) and non-core enriched (gray) transcript/proteins. Abbreviations: DDA: Data Dependent Acquisition; DIA: Data Independent Acquisition; ER: Estrogen Receptor; FDR: False Discovery Rate; GSEA: Gene Set Enrichment Analysis.

Conversely, the most enriched pathways in spiculated tumors included the complement (high enrichment at the protein level) and the EMT gene sets (**Fig 4B-C**), in which a high number of extracellular proteins were found enriched. In the complement pathway, innate immunity response molecules, proteins involved in lipoprotein metabolism (e.g. APOC1, CD36), extracellular matrix proteins (e.g. SERPINA1), and coagulation factors (e.g. F2) were found enriched, though their expression was markedly different from their transcript counterparts. The complement pathway is among the pathways with the highest proportion of secreted proteins that typically operates in the blood plasma and the extracellular space to fight microbes and to remove apoptotic cells, generally as a consequence of actual protein secretion from cells in circulation or in an area surrounding the the tumor. While the complement and EMT pathways include several extracellular and secreted proteins (**Fig S8A**), of which some are shared between the two networks (e.g. FN1), molecules in the EMT set displayed concordant enrichments at the transcript and protein levels. Here several extracellular matrix proteins involved in tissue remodeling (e.g. MMP2, COL1A2) showed comparable RNA and protein levels (**Fig 4C**). This difference may stem from the fact that while complement proteins are secreted by other cell types and distribute within the body through blood vessels, matrix remodeling proteins are generally secreted by the tumor itself or by the surrounding tissue (e.g. fibroblasts). This was implicitly confirmed by the shift in abundance and enrichment score distributions of secreted proteins observed at the MS level within the complement pathway (**Fig S8B**). Conversely such a shift was not observed for the EMT pathway. These data corroborate the notion that proteins expressed in circulation or generally secreted by cells distal to the tumor cannot be accurately evaluated by RNA expression analysis, and that these should be complemented by protein ones.

Overall, the enrichment of networks with a high component of secreted proteins in spiculated tumors suggests a marked interaction between the cancer mass and its surrounding tissue. Proteins dedicated to extracellular matrix remodeling (e.g. MMP2) and organization (e.g. FBLN5), as well as molecules involved in the induction of a mesenchymal cell state (e.g. VIM, FN1; **Fig 4C**) were found enriched both at the RNA and protein level. This suggests reprogramming of the tumor front for tissue invasion. Given the fact that *spiculae* protruding from tumor masses are signs of tumor spread into the surrounding normal tissue, it is likely that remodeling of the extracellular matrix takes place in such tumors. Our data confirms the combined proteogenomics data set can recapture key biological features of breast tumors at the proteome and transcriptome levels, such as the increased immunogenicity and genomic instability of ER negative tumors (Koboldt *et al*, 2012), as well as clarifying the discrepancy between RNA and protein level analyses. In addition, we demonstrate that this strategy can shed light on so far uncharacterized cellular dynamics underlying spiculated cancer appearance, where the stroma is re-arranged around the tumor mass to facilitate invasion. Conversely, the processes operating in non-spiculated tumors seems to revolve around cell proliferation pathways, thus indicating that appearance features might be related to different cell fates (e.g. proliferation vs invasion).

### Evaluation of proteogenomics features

As outlined above, the proposed workflow facilitates proteogenomics analysis of DTU and SNV/SAV expression. For DTU analysis the BANDITs workflow was employed to define differentially expressed transcripts belonging to the same gene in our RNA dataset. These features were then translated at the library level by integration into our spectral library generation and search workflow (**Fig S4** and *Materials and Methods*). The analysis between ER positive and negative tumors generated a total of 539 significant cases of differential transcript usages belonging to 451 genes (FDR cutoff: 0.03). However, pathway enrichment analysis of the 451 genes showing evidence of DTU in the ER+/− comparison revealed no significantly enriched pathway (using the ReactomePA R/Bioconductor package in version 1.30.0; (Yu&He, 2016)).

The same analysis was performed for the appearance group comparison (spiculated vs. non-spiculated) which detected 63 differentially used transcripts in 55 genes (at FDR 0.03). Here the ReactomePA pathway enrichment analysis returned only one enriched pathway: “Endosomal/Vacuolar pathway” (q-value = 0.032), which relates to MHC-1 antigen presentation and adaptive immunity. However, given the relatively sparse input data (only 55 genes were considered, see above) this result needs to be considered carefully and functional assays would be required to confirm the role of the adaptive immune system in its relation to tumor appearance.

Out of the 539 differentially used transcripts in the ER+/− comparison, 127 were subsequently detectable at the proteomic level, and in total 6 were identified by isoform-specific peptides (**Fig 5A**). Interestingly, upon translating our DTU findings onto the protein plane, we noticed that some of the measured variant transcript and protein levels showed differential group enrichment (e.g. ER status: ACAP1, **Fig 5B**; Appearance: DST, **Fig 5C**). In this regard, we assume that the detected discrepancies between transcript and isoform-specific peptide abundances might relate to post-translational regulation of protein abundance, as shown in recent studies (Johansson *et al*, 2019). A similar approach was applied to detect SAVs (see Methods and **Fig S4D**). So far, there has only been a limited number of SAVs that have been confirmed using protein measurement techniques as antibody-based techniques typically are unable to distinguish SAV and MS-based proteomics experiments typically suffer from the limited coverage of peptides per protein. In our workflow, we can partly circumvent this problem by specifically targeting peptides with known SAVs, resulting in the identification of nine high-confident peptides with SAVs (**Fig 5D**). Annotation of the quantified SAVs peptides revealed that several of the peptides that were identified in multiple samples stemmed from variants known to be prevalent in the Nordic population. One SAV-specific peptide however (ATP5PF V69I; **Fig 5E**), was detectable in only one tumor and had no documented prevalence in the Nordic population, indicating a case where a novel mutation detectable at the protein level occurred in cancer.

**Figure 5.**
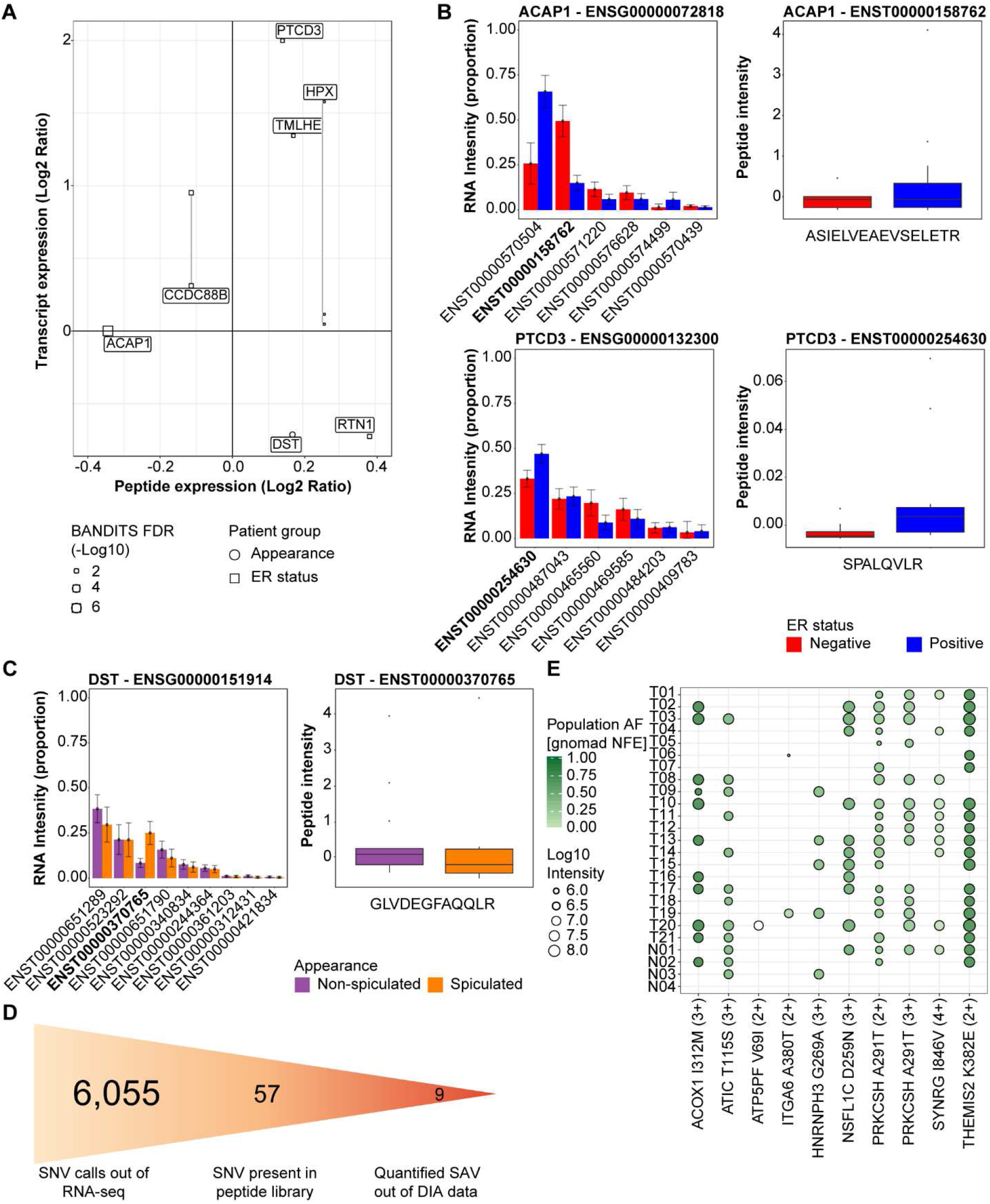
Evaluation of differential transcript usage and Single Amino acid Variant detection at the proteomic level. We employed transcriptomic data information to search our DIA data for DTU (panels A-B) and SAVs (panels C-D). For DTU analysis, we employed the BANDITs workflow to define transcript differential expression, to then generate an isoform-aware spectral library with which to search our DIA MS data. Panel A displays detected DTU at the protein (DIA MS) level and their expression compared to transcript levels. Examples of transcript (left) and (when detected) their specific peptide (right) expression are shown in panel B (ER status) and C (Appearance). For SAV detection, SNVs detected at the RNA level in breast tumors and healthy breast tissue derived from reconstruction surgery were employed to define a variant-specific library against which the DIA data was searched. Panel D displays the number of variant identifications at the RNA and protein level, while panel E shows in which samples (healthy breast tissue and cancer) each variant was detected. Abbreviations: DIA: Data Independent Acquisition; DTU: Differential Transcript Usage; MS: Mass Spectrometry; SAV: Single Aminoacid Variant; SNV: Single Nucleotide Variant.

### Molecular signatures evaluation and drug targeting strategies based on integrated data

In the final analysis, we assessed whether protein-transcripts pairs showed similar trends for key gene signatures currently being explored in routine diagnostic analyses (Mammaprint™, Oncotype-DX™, and PAM50 (Duffy *et al*, 2017)) and targets of FDA approved drugs. In the case of established prognostic signatures such as MammaPrint™, Oncotype-DX™ and the PAM50 classifier, protein-transcript pairs generally displayed high correlation coefficients, as also shown in recent previous studies (Johansson *et al*, 2019). These results suggest a high robustness of these markers and the partial transferability of prognostic signatures from transcriptomics to proteomics (**Fig S9A-B**). On the other hand, correlation of matching RNA-protein pairs for FDA drug targets displayed a considerably wider correlation range (Spearman Rho range: −0.475 to 0.740), suggesting that so far uncharacterized post-transcriptional or translational processes impact on the protein abundances for some of these therapeutic targets. In line with the previously mentioned concept that different mechanisms of transcript and protein regulation might be operating according to tumor context (e.g. ER expression, molecular subtype), we evaluated whether transcript-protein correlations were impacted by receptor status. We verified that several transcript-protein pairs often displayed radically different correlations dependent on ER status, including FDA drug targets such as JAK1 or NFKB1 (**Fig S9C-D**). To further investigate this discrepancy, we extracted co-regulated clusters from the ER positive and negative subsets of our proteomic layers (**Fig S10&Fig S11**). The most represented clusters comprised metabolism, cellular transport, and immune response pathways (**Fig 6 &Fig S12**). Here, even functional protein clusters showed discrepant correlations with RNA data in relation to ER status (e.g. Cell metabolism, **Fig 6A-B** and **Fig S12A-B**). In addition to this, RNA-protein correlations of FDA approved targets such as C1R, DNMT1, KCNQ1 and PRDX5 displayed different RNA-protein correlation coefficients (**Table S6-7**; examples are displayed in **Fig 6C-D** and **Fig S12C-D**). These results confirm that the same gene-protein pair or pathway could be subjected to post-translational regulation mechanisms, but also rely upon different regulatory mechanisms depending on the overall clinical and pathological features of the tumor. While ER status and type of measurement platform have an effect on RNA/protein abundance is not surprising, this phenomenon might have a significant impact in the downstream evaluation of potential biomarker candidates, as differences in transcript-protein correlations are related to patient survival (**Fig 6E** and **Fig S12E**). In the light of this, potential prognostic markers should be either chosen based on RNA or protein data only, taking into account key tumor subgroups/features and, as previously mentioned, the extent of post-translational regulation or class each protein relates to.

**Figure 6.**
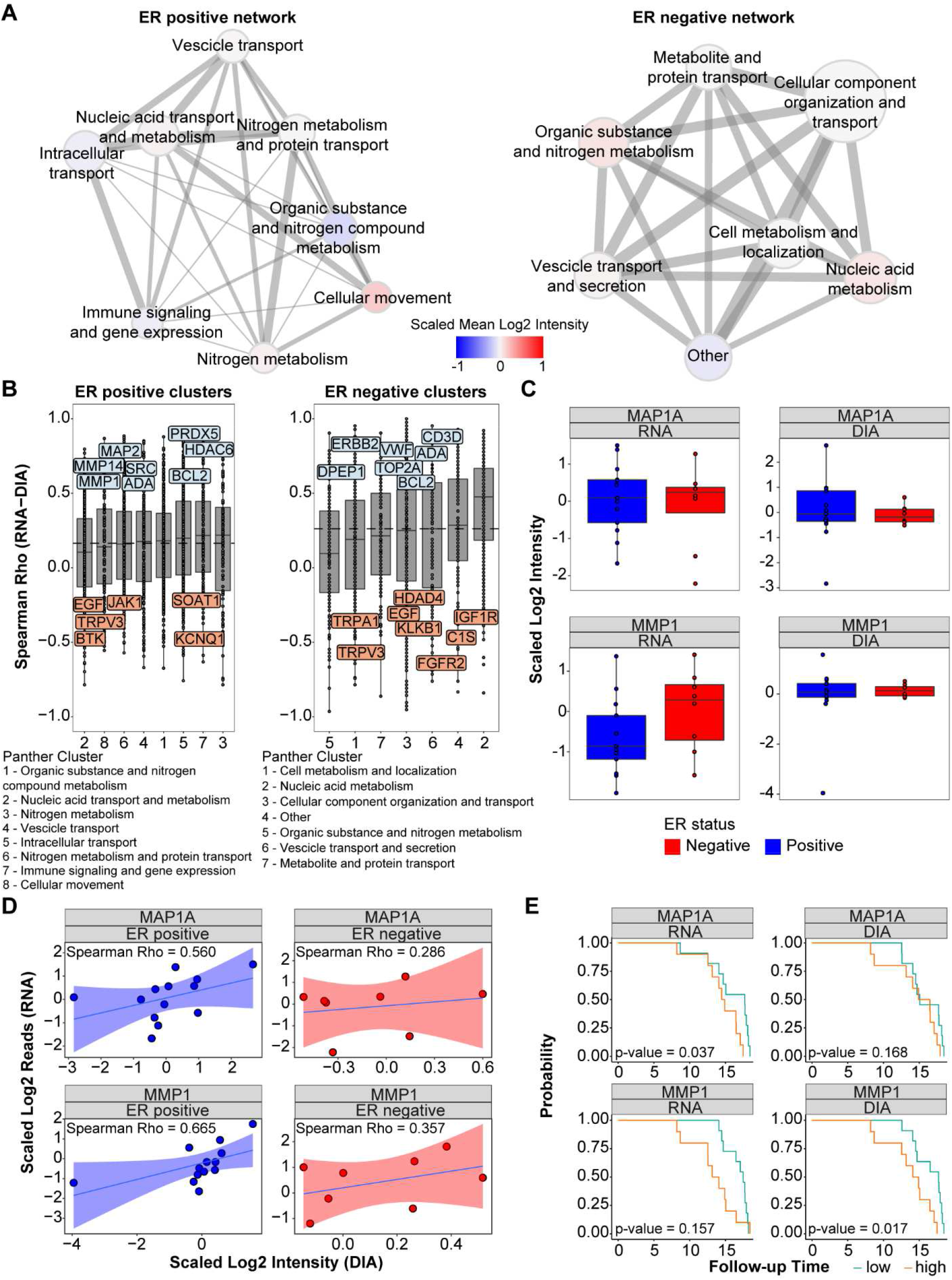
Protein cluster regulation dependent on Estrogen Receptor status. Co-regulated protein clusters in ER positive (left) and negative (right) tumors (see Fig S15) were extracted, annotated with GOBP terms, condensed, and visualized in Cytoscape (panel A). Edge thickness and length relates to cluster distance (Euclidean), node color relates to the scaled mean intensity of all protein in each cluster, and node size depends on the number of proteins in each cluster. Panel B shows the correlation to mRNA of each protein per cluster for ER positive and negative tumors. Marked proteins represent FDA approved drug targets. Comparison of RNA and MS (DIA) expression between ER positive/negative tumors, correlation between RNA and DIA data points in each ER status subgroup, and Kaplan-Meier plots based on RNA and DIA expression (cutoff: median value) are shown if panel C, D, and E, respectively. Abbreviations: DIA: Data Independent Acquisition; ER: Estrogen Receptor; GOBP: Gene Ontology Biological Process; MS: Mass Spectrometry.

Collectively, these results indicate that differentially regulated protein networks exist in clinically relevant sample groups depending on ER status (and possibly mammographic features) and that these protein networks impact both cancer biology as well as the abundance of potential biomarkers and drug targets. While the evaluation of new biomarkers might generally be restricted to the nucleic acid or protein analysis, drug treatments affect proteins in cells, therefore a protein-wide evaluation of these targets is of considerable relevance. Furthermore, to better understand the molecular mechanisms regulating transcript and protein abundance, integrated studies followed by functional assays should be the method of choice to shed light on the fine regulation of key cancer genes and their protein products.

## Discussion

BC is the most common malignancy in women, though its death rate continues to decline due to constant advancements in clinical care, drug target development, and better definition of tumor biology (DeSantis *et al*, 2019). Several key mechanisms underlying breast cancer biology have been elucidated in detail over the years, such as the immunogenicity of triple negative tumors, or the action of the ER transcription factor in ER positive cancers (Prat&Perou, 2011). Despite of this, the mechanisms underlying breast cancer therapy resistance (e.g. ESR1 mutations (Toy *et al*, 2013)), or other prognostic factors such as appearance (e.g. spiculated appearances (Sartor *et al*, 2015)) have yet to be thoroughly investigated.

In BC, recent studies have shown that the integration of genomic and proteomic approaches expanded the knowledge in biological networks underlined by the intrinsic molecular subtypes, and suggested that discrepancies between RNA and quantitative protein data might derive from RNA and/or protein regulation mechanisms (Johansson *et al*, 2019; Mertins *et al*, 2016). Here we developed four improvements of a data independent acquisition mass spectrometry proteogenomics workflow to achieve high proteome depth and quantitative accuracy out of single shot analyses, compare transcriptome data to protein analyses, assess the capability of DIA MS in detecting genomic features such as DTU and SAVs, and to investigate biological pathways underlying understudied tumor appearance features. The combined output from the workflows improved the identification rates from the DIA-MS data. The spectral library used for DIA-MS data analysis was based on DDA MS runs for 21 breast cancer tissues and 4 normal breast specimens, which was further extended with RNA guided assays for SAVs and DTU. The number of identifications we obtained through DIA MS was similar to the ones achieved through fractionated DDA analysis of the same samples, though with fewer missing observations. Additionally, targeted analysis of changes in the mutational and splicing landscapes using informed spectral assays enabled quantification of SAV and DTU-specific peptides. Although the number of quantified SAVs and DTUs was relatively sparse, the possibility of using genomics-informed assumptions to identify the translation of splicing and mutational events at the protein level opens up interesting possibilities for future studies. Further improvements to the computational workflows and data acquisition strategies are required to yield a higher number of identifications out of our DIA data, which can be recurrently mined.

Upon comparing the dynamic range of our transcriptomic and proteomic datasets, we noticed that most transcript-protein pairs displayed similar quantitative levels across our dataset, as observed in previous studies (Wang *et al*, 2019). Only a small subset of proteins displayed negative correlation coefficients with their matching transcript abundances, confirming findings from previous reports (Johansson *et al*, 2019). Here we discovered that proteins involved in processes such as splicing and translational regulation tend to correlate poorly with their transcripts, as opposed to those belonging to immune-related pathways. In this case, cellular processes controlling cellular turnover such as ubiquitination and proteasomal degradation (Kim *et al*, 2011), miRNA activity (Mogilyansky *et al*, 2016; Lorenzo-Martín *et al*, 2019), or epigenetic factors, may actually be responsible for these quantitative discrepancies and target specific protein clusters. The relatively low number of negatively correlated protein-transcript pairs suggest that post-translational regulation might indeed target specific protein groups, but that the impact on the entire proteome might not be as extensive as previously thought (Eraslan *et al*, 2019). For this, further studies of functional nature are required to verify such claims.

Following our analysis of enriched gene-protein pairs and pathways expressed according to the expression of key transcription factors (ER status) or tumor appearances (spiculation) we noticed that RNA and MS measurements converged at the pathway level. This was especially true for previously characterized breast cancer pathways, such as the enrichment of ER responsive genes or immune signaling molecules in ER positive and negative tumors, respectively. In addition to this, our analyses elucidated relevant molecular differences between spiculated and non-spiculated appearances, where tissue remodeling and EMT pathways were found enriched in the former and inflammation- and proliferation-related networks were enriched in the latter. These results confirm that breast cancer invasion of the surrounding tissue through spiculation has been generally associated with stromal and extracellular matrix remodeling. The results also imply that a possible different transcriptional program takes place in these cancers. While EMT has been reasoned to be a mutable transcriptional program (Dongre&Weinberg, 2019), with cells acquiring a spectrum of biological features related to epithelial or mesenchymal fates, our data indicates that invasion of normal tissue through *spiculae* might rely on a mesenchymal cancer cell front. However, future studies are necessary to confirm these results through for example mechanistic experiments such as overexpression studies of the molecular mechanisms related to EMT activation and to establish the association between these features and patient prognosis.

Based on the results that transcriptomic and proteomic analyses largely converge at the pathway level, we further investigated if this also holds true for biomarkers or drug targets. Interestingly, we observe that transcript-protein pairs belonging to established predictive signatures (e.g. Mammaprint) display a high level of correlation, thus suggesting the transferability of these biomarker panels onto the proteomic level. In contrast, this did not hold true for FDA approved drug targets. Since previous studies have shown that post-translational regulation mechanisms might significantly impact protein abundances of drug targets (Johansson *et al*, 2019), we hypothesized that such mechanisms might be operating at different activity levels within critical pathological subgroups such as ER positive and ER negative tumors. Overall, we note that FDA approved drug targets displayed variable degrees of concordance between the two data layers, with foreseeable repercussions in biomarker identification and monitoring dependent on the measurement technology as well as tumor subgroup inherent biology. The expression of differential or mutually exclusive transcriptional programs or regulatory mechanisms is a known factor in cancer (Fujita&Nonomura, 2019). Such transcriptional programs constitute a factor for tumor diversity that establishes cellular changes through genetic and epigenetic mechanisms (Carroll *et al*, 2016; Mohammed *et al*, 2015). These mechanisms may indeed affect genes and proteins on different levels to alter their expression. For this, we find it to be imperative to overlay genomic and proteomic information in order to (i) determine disease subgroups with altered gene and protein expression clusters and (ii) use such information to derive tumor proteotype-specific biomarkers or alternative drug targets and (iii) choose the most appropriate treatment strategy based on tumor subgroup. These findings indicate that the evaluation of protein levels should be performed for a subset of the proteome when evaluating the association of potential markers in the clinical laboratory, or when using mRNA as a substitute for protein abundance. These results support the complementarity of genomic and proteomic information in the dissection molecular pathologies, such as the definition of pathways of interest for further functional assessment and/or drug testing. It is important however, to point out that this study is based on a relatively small patient sample set, which limits the generalization of significant findings out of our analyses.

In conclusion, we have here established improved DIA MS-based workflows in proteogenomic studies to identify mutational processes at the protein level, and the discrepancies that arise between mRNA and protein quantitative data layers, which are in turn dependent on transcript and protein regulation processes. Our analyses also validated previously established enrichments of estrogen receptor-dependent molecular features relating to transcription factor expression, and provided evidence for molecular differences related to the development of mammographic morphologies in spiculated tumor masses. These results demonstrate that there are differentially regulated protein networks in clinically relevant sample groups, and that these protein networks impact both cancer biology as well as the abundance of potential biomarkers and drug target abundance. To assess whether these findings related to biological regulation of protein stability or mRNA translation rates, biochemical and genetic/epigenetic studies should be performed by for example functional high-throughput knockdown models.

## Materials and Methods

### Patients

Out of a large breast cancer dataset consisting of 172 samples, a subset of 21 frozen breast tissue specimens was collected for this study (**Fig 1A**) (Sjöström *et al*, 2018). All specimens were collected from women with primary breast cancer, who underwent tumor resection between 1991 and 2004. Estrogen and Progesterone receptor (ER, PgR) statuses were assessed in tumor tissues by quantitative biochemical assays. Tumor content was derived from microscopic analysis of tumor cell area in hematoxylin-eosin stained tissue slices by two independent trained researchers.

The most dominant mammographic appearance of the tumor was retrospectively collected by one specialist in radiology (HS). Tumors were categorized as spiculated or other tumor appearances such as micro-calcifications or well-defined masses (non-spiculated) based on their most dominating mammographic feature (i.e. appearance categorization). Overlap between spiculation and ER statuses is displayed in **Fig 1B**. Examples of mammographic images are shown in **Fig 1C**). An additional set of 4 normal breast tissues was also collected after breast reduction surgery at Lund University Hospital (SK), and used to generate the DIA spectral library. All tissues were collected from Lund University Hospital and affiliated clinics located in the Skåne region. Clinical and histopathological characteristics of breast cancer patient derived specimens are reported in **Table S1**.

This study used primary breast tumor tissues under approval from the Ethical Review Board (Etikprövningsnämnden) with number DNR 2010/127.

### RNA and Protein Extraction

All breast cancer specimens (n = 21) were processed through AllPrep (Qiagen) protocol for the lysis and extraction of RNA and proteins (**Fig 1D**). Except for the extraction of total protein content, all protocols were performed according to manufacturer instructions (AllPrep RNA extraction kit). An amount of 20-30mg of frozen tissue were cut and collected into tubes for downstream RNA and protein extraction. An adjacent piece, or imprint in cases where not enough tissue for embedding was available, was taken for microscopy and evaluation of cancer content at the center performing RNA extraction. Steel beads (ID 79656, Qiagen) were added to each sample tube together with 400μL a 1% β-mercaptoethanol in RLT buffer (Qiagen) and 2μL of antifoam agent (ID 19088, Qiagen). Tissue disruption was then performed in a Tissue Lyser LT (Qiagen) for 4 min at 50Hz, after which a second volume of 400μL a 1% β-mercaptoethanol in RLT buffer (AllPrep DNA/RNA Minikit, Qiagen) was added after steel bead removal. Samples were then centrifuged at 14,000xg for 5 min. Supernatant was transferred to a new tube kept at −80ºC until RNA and protein extraction.

RNA extraction was performed as per manufacturer instructions using the AllPrep RNA Minikit (Qiagen). Flow-through of each column constituted the protein fraction, which was collected and stored at −80ºC prior to MS sample preparation.

### RNA Quality Control and sequencing analysis

The amount, concentration and quality of the extracted RNA was tested using a Bioanalyzer 2100 instrument (Agilent Technologies, CA, USA), NanoDrop ND-1000 spectrophotometer (Thermo Fisher Scientific, MA, USA) or Caliper HT RNA LabChip (Perkin Elmer, MA, USA). All samples had a RNA integrity value (RIN) of > 6.0.

RNA sequencing analysis was conducted as previously described (Svensson *et al*, 2017). Briefly, RNA concentration was measured in all AllPrep RNA eluates using a Qubit fluorometer following preparation with Qubit RNA HS Assay (Thermo-Fisher). A total of 100 ng of RNA input was then used for cDNA library preparation using the TruSeq® Stranded mRNA NeoPrep kit (Illumina), according to manufacturer instructions. Library cDNA concentration was then measured using the QuantIT® dsDNA HS Assay Kit (Thermo-Fisher), according to manufacturer instructions. cDNA libraries were then denatured and diluted according to the NextSeq® 500 System Guide (Protocol #15048776; Illumina). RNA sequencing analysis was then performed on a NextSeq 500 (Illumina) sequencer generating paired-end reads of length 77bp.

### Protein quantitation and digestion

Collected protein flow-throughs (FT) were subjected to protein precipitation for downstream protein content determination using bicinchoninic acid assay (BCA, Thermo Fisher) assay and trypsin protein digestion. Protein extraction was performed by collecting the flow-through of RNeasy spin columns and by performing protein precipitation as follows: for every sample tube, 3 volumes of acetone were added and samples were incubated at −20ºC for 1 h. Samples were centrifuged at 14,000xg for 20 min and supernatant was removed. Protein pellets were washed twice with 200 μL of 95% ethanol solution for 10 min at room temperature. For FT samples, protein quantitation was performed by BCA assay after re-suspension of the protein pellet in 1X PBS.

For whole tissue lysate (WTL) preparation, frozen tissues were prepared according to previously published protocols (De Marchi *et al*, 2016b),with minor modifications. Briefly, 10 slices of 10 μm thickness were cut for each frozen specimen and re-suspended in ~100μL of ice cold RIPA buffer (150mM sodium chloride, 1.0% NP-40 substitute, 0.5% w/V sodium deoxycholate, 0.1% w/V sodium dodecyl sulphate, 50mM tris-(hydroxymethyl)aminomethane, pH 8.0) supplemented with Halt™ Protease Inhibitor Cocktail (Thermo-Fisher), and sonicated in a cooled Bioruptor-type sonicator (Diagenode) for 15 min. Lysates were then centrifuged at 14,000xg for 20 min at 4ºC and supernatants were transferred in a new tube. Protein concentration was measured by BCA assay (Thermo Fisher).

For downstream trypsin digestion, proteins were precipitated in acetone and washed in ethanol solution (as previously described (Wiśniewski *et al*, 2009)) Briefly, precipitated protein pellets were then re-suspended 100mM Tris (pH 8.0) buffer containing 100 mM dithiotrheitol and 4% w/V sodium-dodecyl-sulphate, and incubated at 95ºC for 30 min under mild agitation. Samples were then cooled to room temperature, and diluted in 8M urea in 100 mM Tris (pH 8.0) buffer for downstream protein digestion. Samples were then loaded on 30 KDa molecular filters (Millipore) and centrifuged at 14,000xg for 20 min. Filters with immobilized proteins were then washed with 100 μ L of 8M urea buffer and centrifuged at 14,000xg for 10 min. Filters with immobilized proteins were then incubated with 8 M urea buffer containing 50mM iodoacetamide for 30 min in the dark. Filters were washed twice with 8M urea buffer, followed by to washes with 50mM tri-ethyl-ammonium bicarbonate buffer (pH 8.0). Proteins were then digested with trypsin (enzyme-protein ratio 1:50) at 37ºC for 16 h under agitation (600 RPM). Filters were then centrifuged at 14,000xg for 20 min to retrieve tryptic peptides.

For DDA MS analysis, a total of 50 ug of protein content were digested for each sample, followed by strong anion exchange fractionation following previously described protocols (Tyanova *et al*, 2016a). Briefly, digested peptides were dried and re-suspended in Britton and Robinson Universal Buffer (20 mM phosphoric acid, 20 mM boric acid and 20 mM acetic acid in ultrapure water; BRUB) pH 11 and loaded on SAX (6 stacked layers; 66888-U, Sigma) stage tips. SAX filters-containing tips were put on top of C18 (3 stacked layers; 66883-U, Sigma) stage tips and peptides were eluted with 100 μL of pH 11 BRUB buffer. SAX stage tips were then transferred onto new C18 tips and peptides were eluted serially at different pH: 8, 6, 5, 4, and 3. C18 tips were then collected, washed with 0.1% formic acid (FA) in ultrapure water and eluted with 100 μL of a solution containing 0.1% FA and 80% acetonitrile in ultrapure water. In order to eliminate any possible remaining contaminants, eluates were dried and subjected to SP3 peptide purification (as described in Hughes *et al* (Hughes *et al*, 2014)). Briefly, 2 μL of SP3 beads (1:1 ratio of Sera Mag A and Sera Mag B re-suspended in ultrapure water; Sigma) was added to dried peptides and incubated for 2 min under gentle agitation. A volume of 200 μL of acetonitrile was then added and samples were incubated for 10 min under agitation. Sample vials were then placed on a magnetic rack and washed again with acetonitrile for 10 min. Elution was performed by adding 200 μL of 2% dimethyl sulfoxide in water to the beads-peptides mixture and incubating them for 5 min under agitation. Supernatant were then collected, dried, and stored at −80ºC until MS analysis.

For downstream DIA MS analysis, a total of 10 μg of protein was digested as previously mentioned, omitting SAX stage tip-based fractionation. Solid phase extraction was performed using the SP3 method, as aforementioned.

### Mass Spectrometry Analysis

Global proteome MS analysis was performed on a Q-Exactive Plus (Thermo-Fisher) mass spectrometer. Around 1 μg of Tryptic peptides from fractionated samples were separated on a RP-HPLC EasySpray column (ID 75 μm × 25 cm C18 2 μm 100 Å resin; Thermo-Fisher) coupled to an EASY-nLC 1000 liquid chromatography system (Thermo Fisher).

For DDA analysis of SAX-fractionated samples, peptides from each fraction (n of fractions: 6) were eluted in a 90 min gradient (flow: 300 nl/min; mobile phase A: 0.1% formic acid in H2O; mobile phase B: 99.9% acetonitrile and 0.1% formic acid). Chromatographic gradient was run as follows: 5% B for 5 min; 5-30% B in 85 min; 95% B for 10 min. The 15 most abundant peaks from the MS scan (resolution: 70,000 at 200 m/z) were selected and fragmented by higher energy induced collision dissociation (HCD; collision energy: 30). Dynamic exclusion was enabled (window: 20 s). AGC target for both full MS and MS/MS scans was set to 1e6. Precursor ions with intensity above 1.7e4 were selected for MS/MS scan triggering.

For DIA MS analysis, unfractionated samples were eluted in a 120 min gradient (flow: 300 nl/min; mobile phase A: 0.1% formic acid in H2O; mobile phase B: 80.0% acetonitrile and 0.1% formic acid) on a Q-Exactive HFX (Thermo-Fisher) instrument coupled online to an EASY-nLC 1200 system (Thermo-Fisher). Digested peptides were separated by RP-HPLC (ID 75 μm × 50 cm C18 2 μm 100 Å resin; Thermo-Fisher). Gradient was run as follows: 10-30% B in 90 min; 30-45% B in 20 min; 45-90% B in 30 s, and 90% B for 9 min. One high resolution MS scan (resolution: 60,000 at 200 m/z) was performed and followed by a set of 32 DIA MS cycles with variable isolation windows (resolution: 30,000 at 200 m/z; isolation windows: 13, 14, 15, 16, 17, 18, 20, 22, 23, 25, 29, 37, 45, 51, 66, 132 m/z; overlap between windows: 0.5 m/z). Ions within each window were fragmented by HCD (collision energy: 30). AGC target for MS scans was set to 1e6 for MS and MS/MS scans, with ion accumulation time set to 100 ms and 120 ms for MS and MS/MS, respectively.

### DDA MS data processing

DDA-derived RAW files were analyzed using MaxQuant (v1.6.0.16). MS spectra were searched using the Andromeda built-in search engine against the Uniprot-Swissprot human proteome database (version download 2017.06.12). LFQ and match between runs options were enabled. Proteins were then filtered for false discovery rate (Q-value < 1%), reverse sequences (excluded), contaminants (excluded), and identification of unique peptides (at least one unique peptide per protein). LFQ intensities were then Log2 transformed and protein-level scaled prior statistical analysis.

### DIA MS data processing and spectral Library Generation (MakeGTL workflow)

The workflow used in the spectral library generation for the DIA search is described in **Fig S3A** and **Fig S4A**. This computational pipeline uses the DDA raw data by first employing MaRaCluster (v0.05.0, Build Date Apr 16 2018 21:04:32) to cluster the spectra, and then generating consensus spectra from the clustering using a 5% clustering p-value cut-off. Other parameters of MaRaCluster were kept to default settings. In the next step, the consensus spectra of the selected clusters were searched against the sequence database (Uniprot-Swissprot version download 2017.06.12 for general protein quantification) using Comet (v”2017.01 rev. 4”; (Eng *et al*, 2013)) and the resulting Peptide Spectral Matches (PSMs) were scored by Percolator (v3.02.1, Build Date Aug 13 2018 15:50:58; (The *et al*, 2016)). The resulting scored PSMs were then processed using an in-house built python script, that selected (for each peptide) the spectral match with the best Q-value smaller than 10%. Subsequently, our script extracted transitions from each spectrum by matching peaks to theoretical ion masses within 1ppm (only y and b ions were considered here, since our machine mostly generates such ions). The resulting output consisted of an OpenSWATH compatible Generic Transition List (GTL) in tsv format. We applied this workflow to three sets of raw DDA files: the dataset generated in this study, the Tyanova (Tyanova *et al*, 2016a) dataset, and the Bouchal (Bouchal *et al*, 2019) dataset.

### Iterative RT peptide selection and quantification in DIA analysis (DIAnRT workflow)

To select a set of internal retention time (iRT) peptides (i.e. peptides that are endogenously present in each sample run) OpenSWATH was run without iRT peptide input in order to extract the best candidates to be used as iRT peptides for the next iteration (**Fig S4B**).

We used a python script to extract from the resulting OSW files a set of peptides that were detected closest to their library retention time (at most 10 minutes). These were then additionally filtered based on peak-width (i.e. less than 16.5 seconds at the base), and intensity (i.e. at least 1e5). Peptides detected in less than 20 samples were discarded and the remaining set of peptides was randomly subsampled to no more than 100 peptides per RT bin when splitting the DIA gradient into 20 equally sized RT bins. This set of peptides was then used in the next step to fit a lasso iRT model with OpenSWATH, and the results were again processed to extract the best-fitting, sharp-peaked peptides that were found in many samples. Each set of output files was scored using PyProphet (v2.0.1) by merging a subsample of 5% of PSMs from each sample and scoring the merged dataset (scoring level was MS1-MS2). The resulting model (build on a representative sample comprised of 5% from each individual sample) was then back-propagated to the individual samples and used for their scoring. The so scored samples were then passed to the feature_alignment.py script for TRIC alignment (Röst *et al*, 2016). After 5 iterations the number of proteins identified with this method seemed to stabilize (4219, 4281, 4298, 4302 and 4301 respectively using the library generated from our DDA data), and for this we chose the last iteration results as our dataset for further analysis.

Subsequent to re-quantification, 16,658 peptides covering 4,217 proteins were quantified. These were scaled by per-sample median intensity to account for sample-level differences. A Log2-transformed, mean-centered and standard deviation-scaled version of this matrix was generated both on the peptide level and on the protein level. For protein summarization we first selected for each protein the larges subgroup of at least 3 peptides that had a spearman correlation of at least among them, summed those peptides intensities and applied the Log2 transformation, scaling and centering on this summed intensity. We applied the same workflow to all three generated libraries (our own, Tyanova and Bouchal) individually.

### Combining results from multiple DIA library searches

For general peptide quantification three libraries (Subtypes, Tyanova and Bouchal) were employed in the DIAnRT workflow to create three sets of peptide quantifications, which were combined before PyProphet (Rosenberger *et al*, 2017) scoring. For each peptide in the superset of quantified peptides from all three libraries, p-values according to the following rules:

1. If the peptide is quantified in our own library we used that p-value.
2. If the peptide is quantified in only one library we used that p-value.
3. If the peptide is quantified in both the Tyanova and the Bouchal library but not in our library we used the quantification with the lower p-value.

Peptide quantifications were then combined into a single table and processed with PyProphet for Q-value calculation, followed by feature alignment between DIA runs and re-quantification based on the alignment.

### RNA data processing

The de-multiplexed RNA-Seq reads were aligned to the GRCh38 human reference genome using STAR aligner (v020201) with an overhang value of 75 to match the read-length. Subsequently we employed the standard GATK analysis pipeline including duplicate removal, indel realignment and base quality score recalibration (GATK v3.7-0-gcfedb67).

The resulting bam files were processed using the DESeq2 R/Bioconductor package (version 1.22.2) by first generating per-gene read counts mapping to the GRCh38 GTF file from Ensembl version 95 using the *summarizeOverlaps* function in “Union” mode to count reads that map uniquely to exactly one exon of a gene. Subsequent to discarding genes with no counts in any of the samples, DESeq analysis was performed with the ER status (i.e. ER positive and ER negative) as the explanatory variable in the model followed by Log-Fold-Change shrinkage. A separate DESeq analysis was also performed using spiculation (i.e. spiculated and non-spiculated) status as the explanatory variable.

For DTU detection (computational workflow is shown in **Fig S4C**), the BANDITs (https://doi.org/10.1101/750018) workflow was employed to analyze the RNA-seq data, and to determine a set of genes with differential transcript usage in the comparison of ER positive and ER negative samples. To verify DTU at the proteome level an isoform-aware spectral library was generated from the Ensembl GRCh38 human proteome by *in silico* tryptic digestion of all protein isoforms found in this database, and the determination for each peptide the set of protein isoforms it matched to. For each unique combination of isoforms, all matching peptides of at least length 5 amino acids were concatenated to create a mock protein sequence specific to that combination of isoforms. Using the resulting FASTA file and our DDA data an isoform-aware spectral library was created, where each detectable peptide was matched to a set of protein isoforms identified by their Ensembl protein IDs. The DIAnRT workflow was employed on this library to quantify peptides in an isoform aware fashion from our DIA data (i.e. by re-using the set of RT peptides generated in the iterative DIA quantification process described above). Quantified peptide intensities of those proteins that matched to genes with significant differential transcript usage were overlaid onto those determined by the BANDITS workflow on the RNA-seq data.

For SNV/SAV evaluation (**Fig S4D**), SNV calls were derived out of the aligned RNA-seq reads using the h5vc R/Bioconductor package with the callVariants function, requiring at least 2 reads supporting the variant and at least 10 reads total coverage. We annotated the SNVs using the Ensembl Variant Effect Predictor and filtered the SAVs in order to retain only those events that modify the amino acid sequence of the affected protein.

By using the set of nonsynonymous SAV calls obtained from the RNA-seq data, we generated (for each SAV) its derived protein sequence, and used in-silico digestion to determine the resulting set of tryptic peptides. By discarding all peptides that also arose from the unmodified reference sequence, a set of peptides that specifically identify each SAV was determined (typically only one peptide, except where SAVs generated new tryptic peptides). From these results a FASTA file containing the concatenated peptide sequences that identify each SAV was created and used this as input within the MakeGTL workflow to create a SAV library in order to quantify the SAVs in our DIA data.

### Statistical and Pathway Analyses

In the comparative analysis of the combined WTL and FT cohorts, proteins with less than 30 % missing observations (< 30% missing data) in the DDA set were included. This resulted in a list of 2,608 proteins. Welch-corrected t test was performed to assess significant differences, followed by Benjamini-Hochberg p-value adjustment as multiple test correction. All proteins were annotated with Gene Ontology Biological Process (GOBP) and Gene Ontology Cellular Component (GOCC) terms, and 1D and 2D enrichment analysis were performed in Perseus (v1.5.5.3) (Cox&Mann, 2012; Tyanova *et al*, 2016b).

In our correlation analyses between transcript and protein abundances, we employed Spearman correlation to calculate both the correlation coefficient and p-value. To assess whether specific protein clusters were affected by different mRNA-protein correlation distributions, all proteins were annotated with GOBP terms, the distribution of correlation coefficients of each GOBP annotation was then tested against the background (i.e. all proteins) by t test, followed by Benjaminin-Hochberg p-value adjustement. Selected adjusted p-value cutoff for GOBP annotation was 0.15.

In the analysis for differential pathway enrichment between ER statuses (ER positive vs ER negative) we performed Gene Set Enrichment Analysis (GSEA (Subramanian *et al*, 2005); database: Hallmarks v5.2; permutation type: gene set; scoring: weighted; metric: t test; other parameters were left at default settings) on RNA, DDA (FT subset only), and DIA data layers. Input data tables were filtered as follows: RNA (no filtering), DDA (< 30% missing observations), DIA (no filtering). Enrichment scores of the top50 significant (i.e. by Q-value) pathways were then plotted for each data-layer.

In order to define co-regulated protein clusters in our DDA and DIA datasets, we generated Spearman correlation-based matrices for the ER positive and negative groups. Using the Elbow method the minimum number of clusters was then defined for each ER status sample group and the results from the Panther over-representation test (database: GOBP complete; http://www.pantherdb.org/) were used to annotate each cluster. A second Elbow method step was performed in order to merge highly similar clusters after calculating their similarity by clustering analysis (metric: Euclidean distance complete clustering). Kaplan-Meier curves were built using total follow-up time as continuous variable, while the event variable was set to 1 for all samples (i.e. all patients survived with no recurrence during the follow up period) to demonstrate the potential impact of transcript-protein measurement on survival-type analyses for the evaluation of putative biomarkers. For these proof-of-concept analyses, patients were split into two groups based on median expression of each transcript-gene (low: < median expression; high: ≥ median expression) within each data layer (RNA, DDA, and DIA) and difference in follow-up time was evaluated by Log-rank test.

## Acknowledgements

The authors would like to thank Kristina Lövgren and Sara Baker for sample collection, microscopic evaluation of tumors, and patient clinical database management. We also thank Carlos Alberto Gueto Tettay for implementation of the MakeGTL workflow.

## Author Contributions

TDM, PTP, MS, JM, LM, and EN conceived and designed the study. MS collected and managed the clinical data. SK selected and provided normal breast tissue samples. HS collected all mammographic images of breast tumor samples and analyzed tumor appearances. TDM prepared all samples for downstream MS analysis. PTP and LM created the DIA computational workflows. TDM and PTP performed data analysis. TDM, PTP, LM, JM, and EN wrote the manuscript. All authors critically reviewed the manuscript.

## Data Availability

Transcriptomic data was uploaded to Mendeley Data under DOI:10.17632/wmkb7z7mz4.1. Raw sequence data was not uploaded in accordance with Swedish law, which prevents the upload of person-specific and – identifying information.

DDA and DIA MS data, and their respective search result files, have been deposited to the ProteomeXchange Consortium via the PRIDE partner repository (Vizcaíno *et al*, 2013) with the dataset identifier: PXD018830.

