## Supplementary Figures and Tables for "“A Proteogenomic workflow reveals distinct molecular phenotypes related to breast cancer appearance”": 20200504_MSB_supp-mat.pdf

9. Current address: Department of Radiation Oncology, University of California, San Francisco, 1450 3rd Street, CA 94158, San Francisco, USA.

#### Supplementary Tables

**Table S1. Clinical characteristics of breast cancer cohort.**

| | **Breast cancer tissues (n = 21)** | |
|---|---|---|
| | **N** | **%** |
| **Age*** | | |
| **≤ 55 years** | 8 | 38.1 |
| **> 55 years** | 11 | 52.4 |
| **Tumor size** | | |
| **T1 (≤ 2mm)** | 13 | 61.9 |
| **T2 (2-5mm) + Tx** | 8 | 38.1 |
| **ER** | | |
| **Negative** | 8 | 38.1 |
| **Positive** | 13 | 61.9 |
| **PgR** | | |
| **Negative** | 8 | 38.1 |
| **Positive** | 13 | 61.9 |
| **Appearance** | | |
| **Spiculated** | 7 | 33.3 |
| **Non-spiculated** | 14 | 66.7 |

*Patients with missing characteristics are not reported

Abbreviations: ER, Estrogen Receptor; PgR, Progesterone Receptor.

#### Figure Legends

**Figure S1. Impact of sample preparation method on proteomic identifications.**

Samples for DDA MS-based library generation were prepared as FT and WTL. After MaxQuant search, we compared the number of unique peptides and proteins identified in each subgroup (A) and their overlap (B). Data density for peptides (C) and proteins (D) were then compared and found to be highly similar. On sequence coverage (E) FT and WTL were highly comparable, though their combination showed a slightly better performance.

Abbreviations: DDA: Data Dependent Acquisition; FT: Flowthrough; MS: Mass Spectrometry; WTL: Whole Tissue Lysate.

**Figure S2. Evaluation of quantitative differences and similarities between sample preparation methods.**

Figure displays the results from the analysis of DDA MS data from our FT and WTL samples. Differentially expressed proteins (t test adjusted p-value $< 0.05$) were plotted in a heatmap (A). Panel B displays scatter plots of 2D analysis-derived GOCC and GOBP enriched terms in FT and WTL samples, while panel C shows the results of 1D analyses in the FT and WTL subsets in relation to tumor area content.

Abbreviations: DDA: Data Dependent Acquisition; FT: Flowthrough; MS: Mass Spectrometry; GOBP: Gene Ontology Biological Process; GOCC: Gene Ontology Cellular Component; WTL: Whole Tissue Lysate.

**Figure S3. Scheme of DIA search and overview of separate library search results.**

Panel A displays a schematic overview of DIA library preparation for all three datasets (i.e. this study (De Marchi), Bouchal and Tyanova; upper panel) and finalization of search results (lower panel). Panel B displays identification overlap for peptides (left) and proteins (right) across the three libraries.

Abbreviations: DIA: Data Independent Acquisition.

**Figure S4. Computational workflow generated to analyze the DIA dataset.**

Computational workflows (see Methods for details) were established in this study to generate spectral libraries (A) for DIA data search (B), assess differential transcript usage at the proteomic level (C), and detect and quantify SAVs (D).

Abbreviations: DB: Database; DDA: Data Dependent Acquisition; DIA: Data Independent Acquisition; RT: Retention Time; SAV: Single Aminoacid Variant; SNV: Single Nucleotide Variant; VEP: variant Effect Predictor.

**Figure S5. Evaluation of p-value distribution out of RNA-protein correlation analyses.**

Following our correlation analyses between matching transcripts and proteins (reported in Fig. 2), we evaluated the distribution of the negative and positive correlation coefficients-related p-values. We noticed that the uncorrected p-values from negative Spearman correlations seemed to follow a background-like distribution (A). These p-values became non-significant after Benjamini-Hochberg correction (B), thus suggesting that they might not be statistically meaningful or that the effect size might be too small.

**Figure S6. Analyses between transcript-protein correlations and proteomic features.**

Spearman correlation analysis was between transcript-protein correlation and protein intensity (A: DDA; B: DIA), and sequence coverage (C: DDA; D: DIA), with no strong association found between these variables.

Abbreviations: DDA: Data Dependent Acquisition; DIA: Data Independent Acquisition; MS: Mass Spectrometry.

**Figure S7. Protein-level comparison between transcriptomic and DDA proteomic data in the context of ER status and spiculated appearance.**

The top 100 differentially expressed transcripts (ordered by p-value; RNA dataset) between ER positive/negative (panel A) and spiculated/non-spiculated (panel B) tumors were matched against the DDA proteomic dataset and the Log2 Ratios between groups were compared. In all panels, dot size is relative to adjusted p-value. Transcript-protein pairs that showed similar expression or enrichment between RNA and proteomic data are shown in blue, while discordant enrichments are displayed in red.

Abbreviations: DDA: Data Dependent Acquisition; DIA: Data Independent Acquisition; ER: Estrogen Receptor; FDR: False Discovery Rate.

**Figure S8. Evaluation of secreted protein effect on transcript-protein discrepancies in pathway analysis.**

Figure display the evaluation of secreted proteins presence in HALLMARKS pathways and their expression in Complement pathway. Panel A displays the ranking of HALLMARKS pathways based on the proportion of secreted proteins in each network. The top 5 pathways are labeled. Panel B displays the enrichment score and abundance (Log2Ratio) distributions of transcripts (RNA) and proteins (DDA, DIA) in the Complement and EMT datasets for non-secreted (light red) and secreted (light blue) proteins.

Abbreviations: DDA: Data Dependent Acquisition; DIA: Data Independent Acquisition; EMT: Epithelial Mesenchymal Transition.

**Figure S9. Evaluation of transcript-protein correlation in biomarker signatures and drug targets and the impact of tumor subgroup on RNA-protein correlation.**

Figure show correlation between transcript-protein pairs for biomarker signatures (Mammaprint, Oncotype-DX, and the PAM50 classifier) for both the DDA (A) and DIA (B) data layers. Panels C and D display transcript-protein correlation coefficients between ER positive and negative tumor groups (panel C: DDA; panel D: DIA). FDA drug targets are marked in red.

Abbreviations: DDA: Data Dependent Acquisition; DIA: Data Independent Acquisition; ER: Estrogen Receptor; FDA: Food and Drug Administration.

**Figure S10. Co-regulation clusters in ER positive and negative tumor samples out of the DDA data layer.**

In order to define the most represented pathways in our DDA MS data, we generated Spearman correlation matrices within the ER positive and ER negative tumor groups (A). We then employed the elbow method to define the minimum number of clusters for both sample groups (ER positive: 12 clusters; ER negative: 9 clusters; panel B). Having observed an overlap between Panther-derived enriched terms (over-representation test for GOBP terms) among clusters, we decided to merge highly similar clusters and calculated the distance between them in the ER positive (C) and negative (D) subgroups. The minimum number of clusters was then selected again based on the Elbow method (E).

Abbreviations: DDA: Data Dependent Acquisition; ER: Estrogen Receptor; GOBP: Gene Ontology Biological Process.

**Figure S11. Co-regulation clusters in ER positive and negative tumor samples out of the DIA data layer.** Following our analysis of clusters for DDA data, we generate Spearman correlation matrices within the ER positive and ER negative tumor groups (A) out of the DIA data layer. The elbow method was used to define the minimum number of clusters for both sample groups (ER positive: 13 clusters; ER negative: 10 clusters; panel B). Highly similar clusters were merged after calculating the distances between them in the ER positive (C) and negative (D) subgroups, and by selecting a new minimum number of protein clusters (E).

Abbreviations: DIA: Data Independent Acquisition; ER: Estrogen Receptor; GOBP: Gene Ontology Biological Process.

**Figure S12. Protein cluster regulation dependent on Estrogen Receptor status out of the DDA data layer.** Co-regulated protein clusters in ER positive (left) and negative (right) tumors (see Fig. S8) were extracted, annotated with GOBP terms, condensed, and visualized in Cytoscape (panel A). Edge thickness and length relates to cluster distance (Euclidean), node color relates to the scaled mean intensity of all protein in each cluster, and node size depends on the number of proteins in each cluster.

Panel B shows the correlation to mRNA of each protein per cluster for ER positive and negative tumors. Marked proteins represent FDA approved drug targets. Comparison of RNA and MS (DIA) expression between ER positive/negative tumors, correlation between RNA and DIA data points in each ER status subgroup, and Kaplan-Meier plots based on RNA and DDA expression (cutoff: median value) are shown if panel C, D, and E, respectively.

Abbreviations: DDA: Data Dependent Acquisition; ER: Estrogen Receptor; GOBP: Gene Ontology Biological Process; MS: Mass Spectrometry.

### Figures

**Figure S1. Impact of sample preparation method on proteomic identifications.**

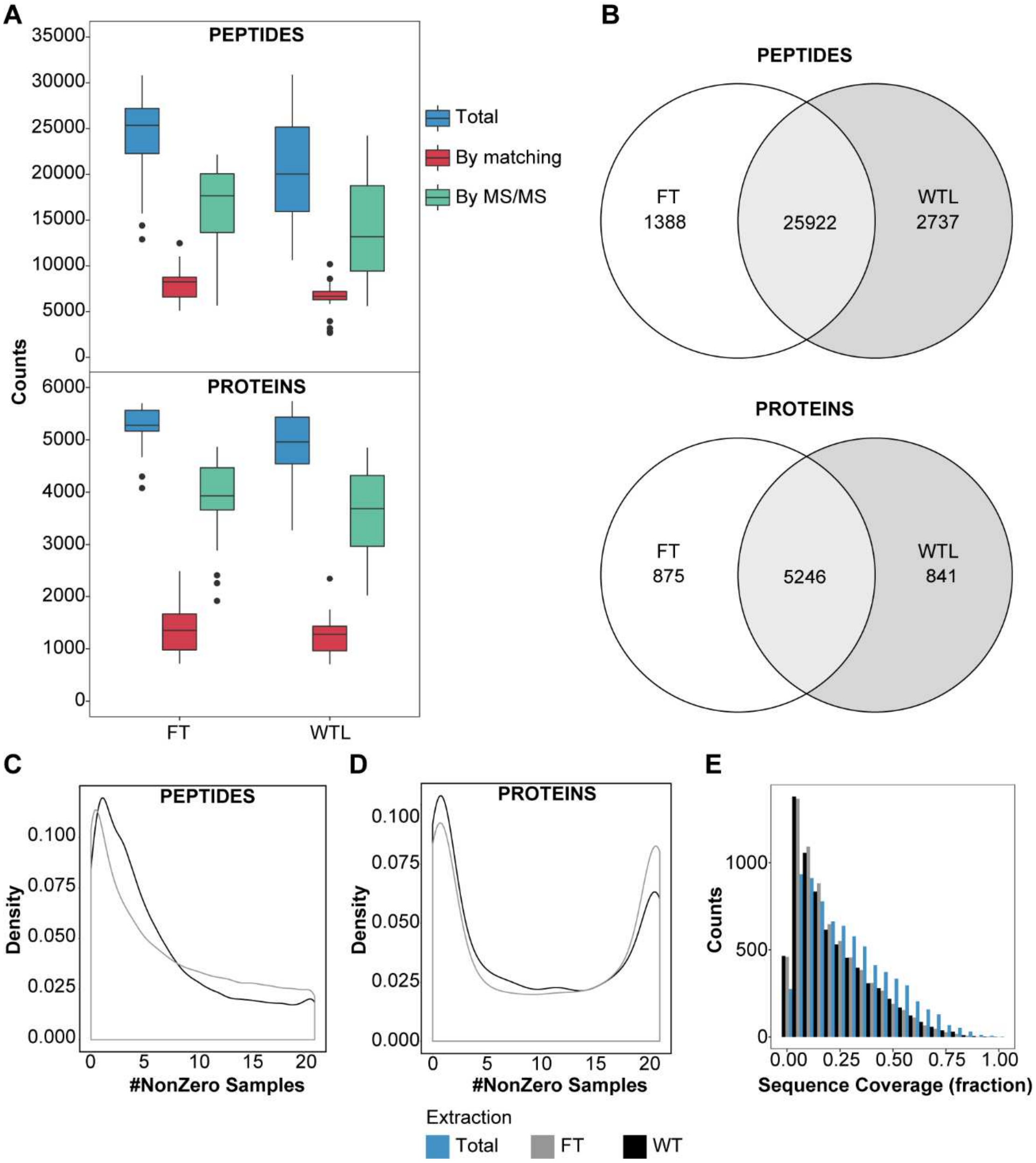

**Figure S2. Evaluation of quantitative differences and similarities between sample preparation methods.**

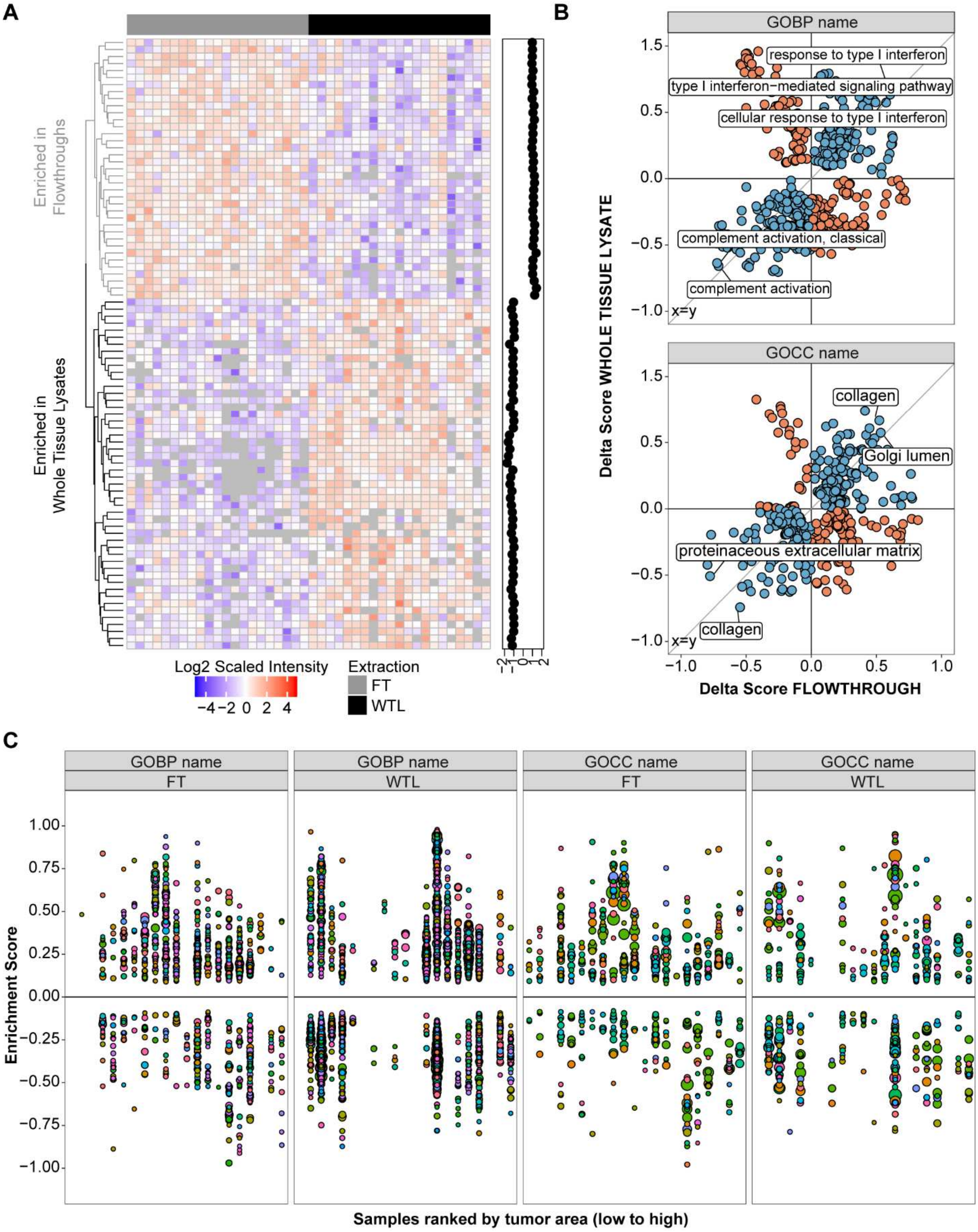

**Figure S3. Scheme of DIA search and overview of separate library search results.**

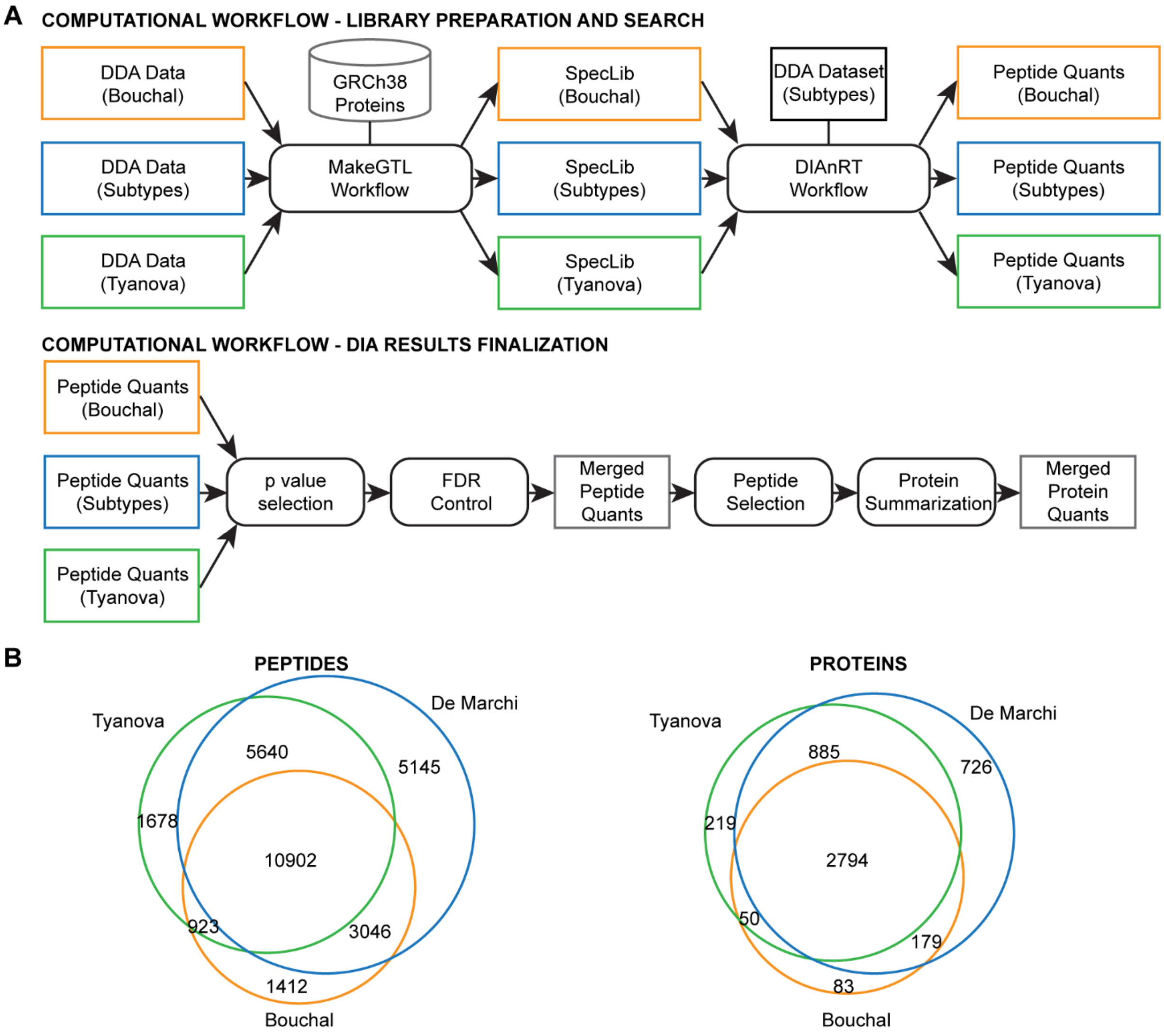

**Figure S4. Computational workflow generated to analyze the DIA dataset.**

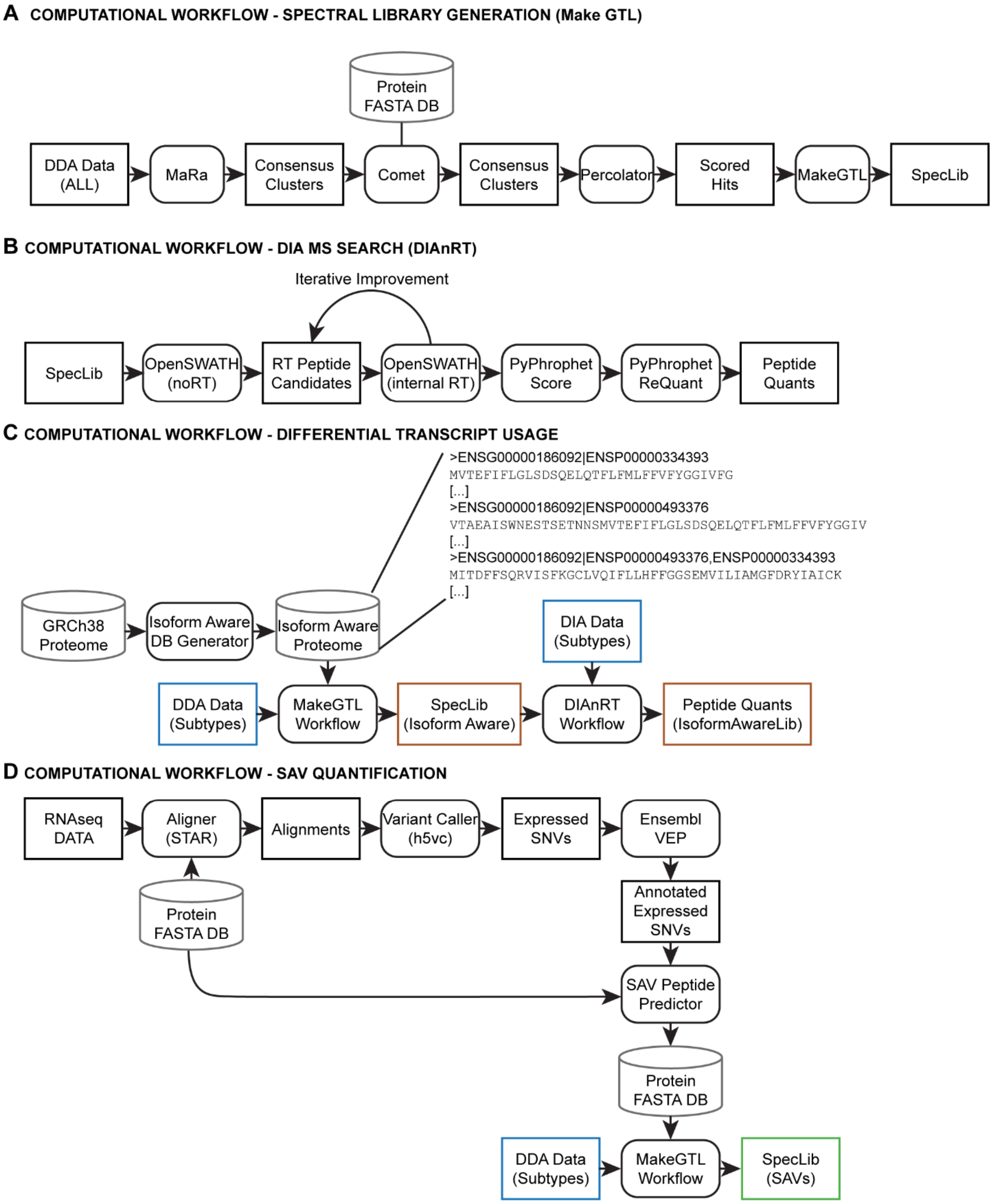

**Figure S5. Evaluation of p-value distribution out of RNA-protein correlation analyses.**

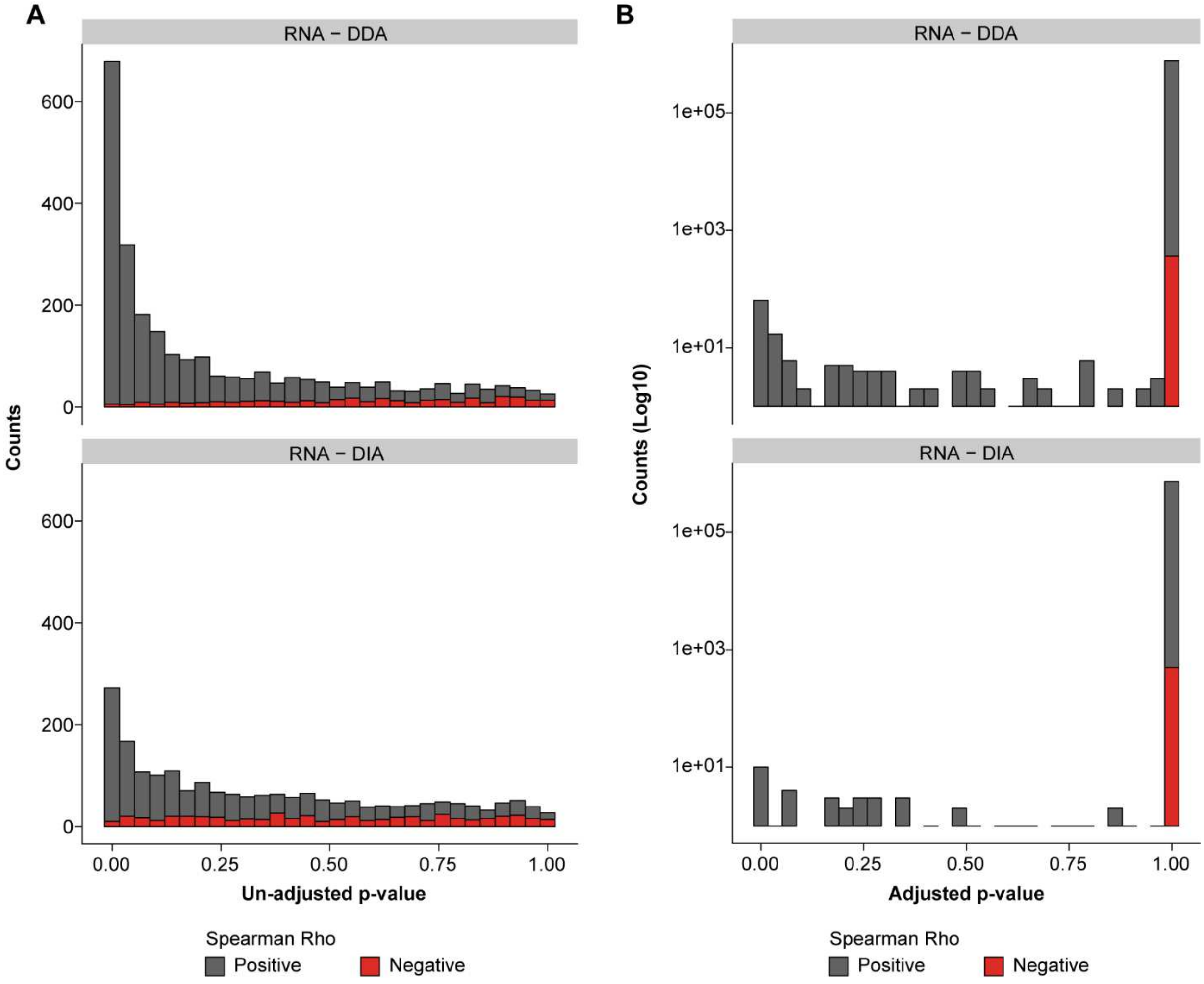

**Figure S6. Analyses between transcript-protein correlations and proteomic features.**

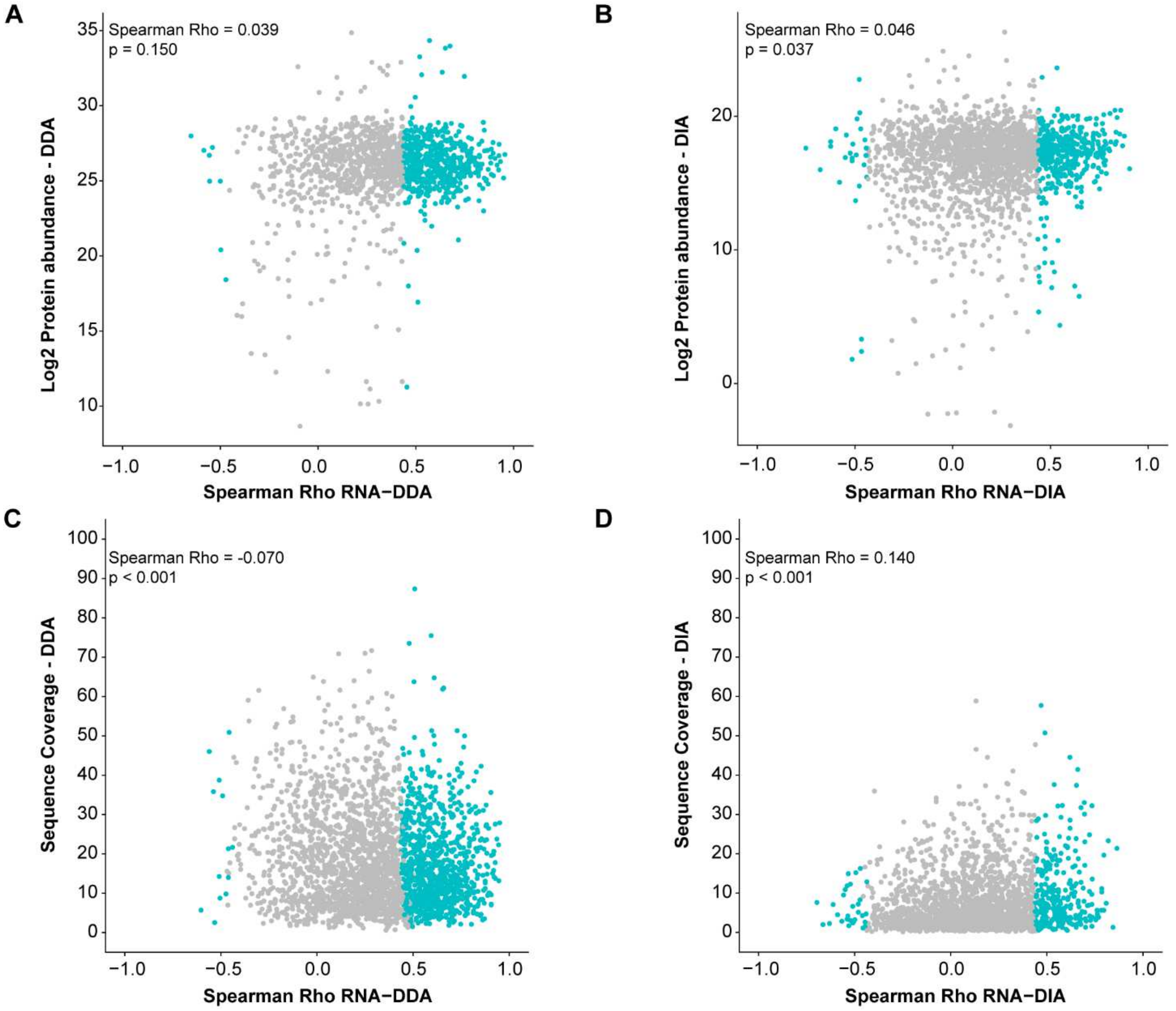

**Figure S7. Protein-level comparison between transcriptomic and DDA proteomic data in the context of ER status and spiculated appearance.**

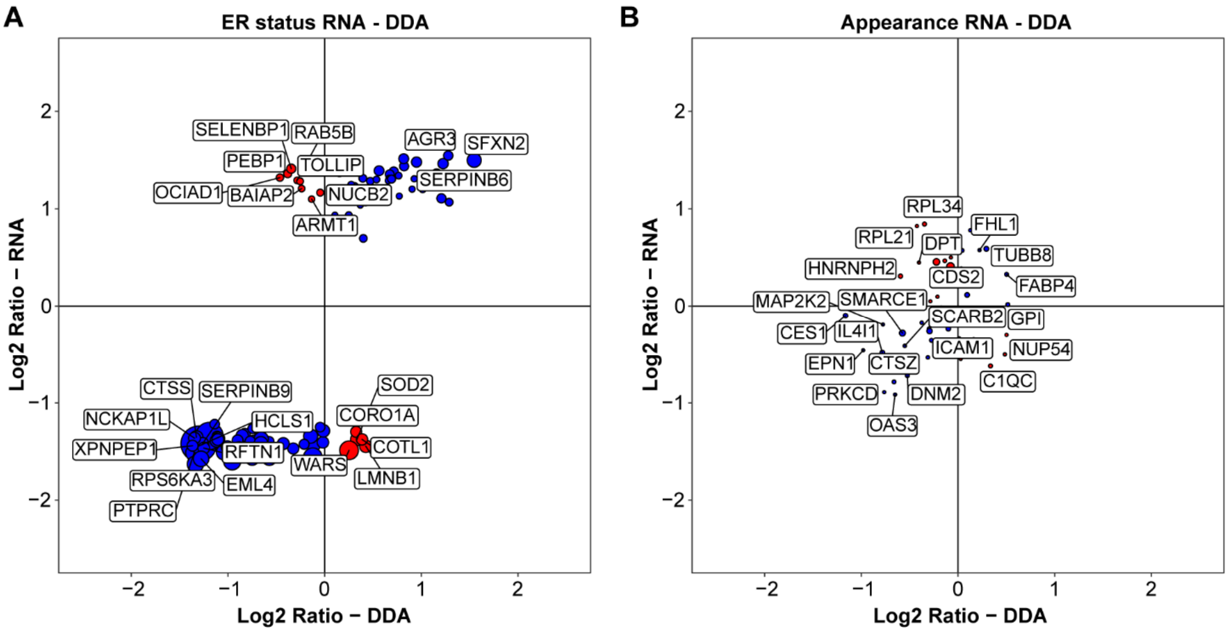

**Figure S8. Evaluation of secreted protein effect on transcript-protein discrepancies in pathway analysis.**

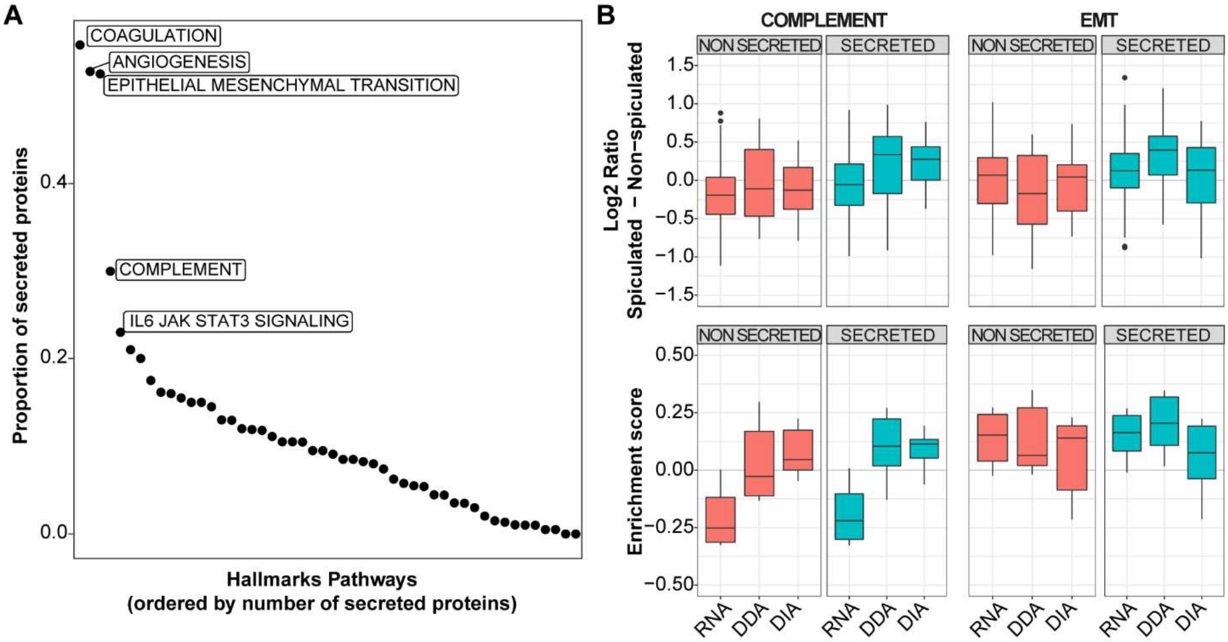

**Figure S9. Evaluation of transcript-protein correlation in biomarker signatures and drug targets and the impact of tumor subgroup on RNA-protein correlation.**

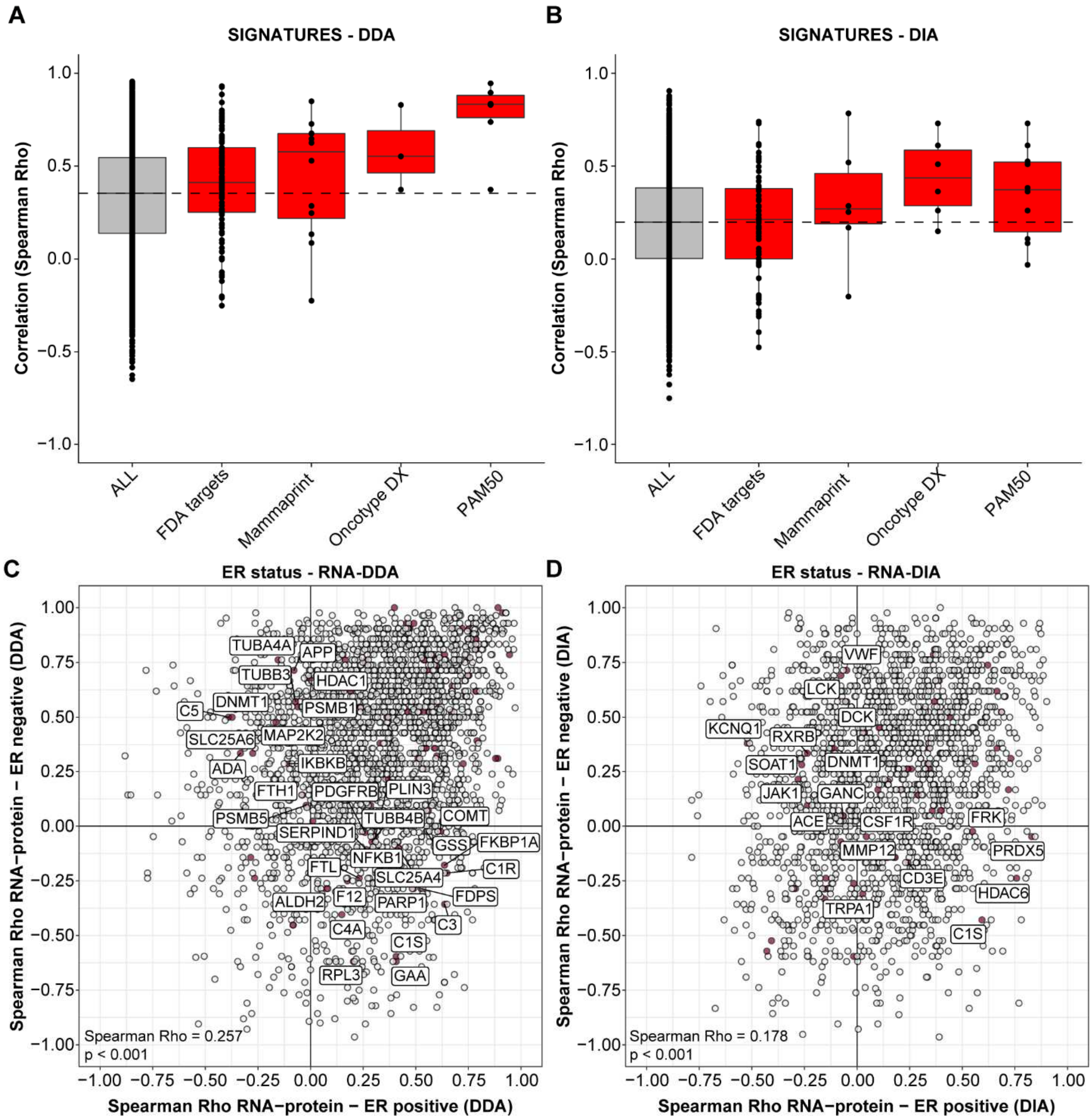

**Figure S10. Co-regulation clusters in ER positive and negative tumor samples out of the DDA data layer.**

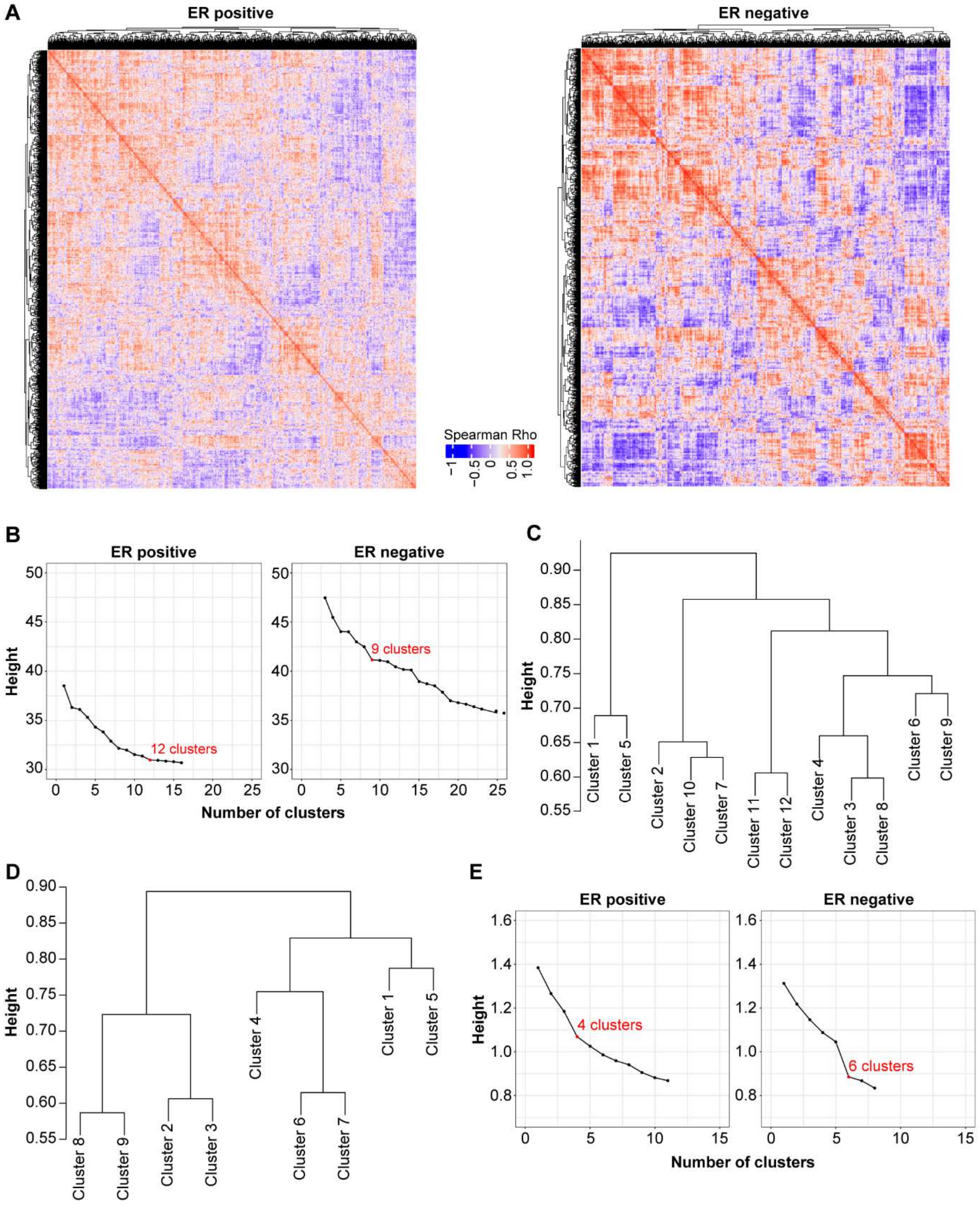

**Figure S11. Co-regulation clusters in ER positive and negative tumor samples out of the DIA data layer.**

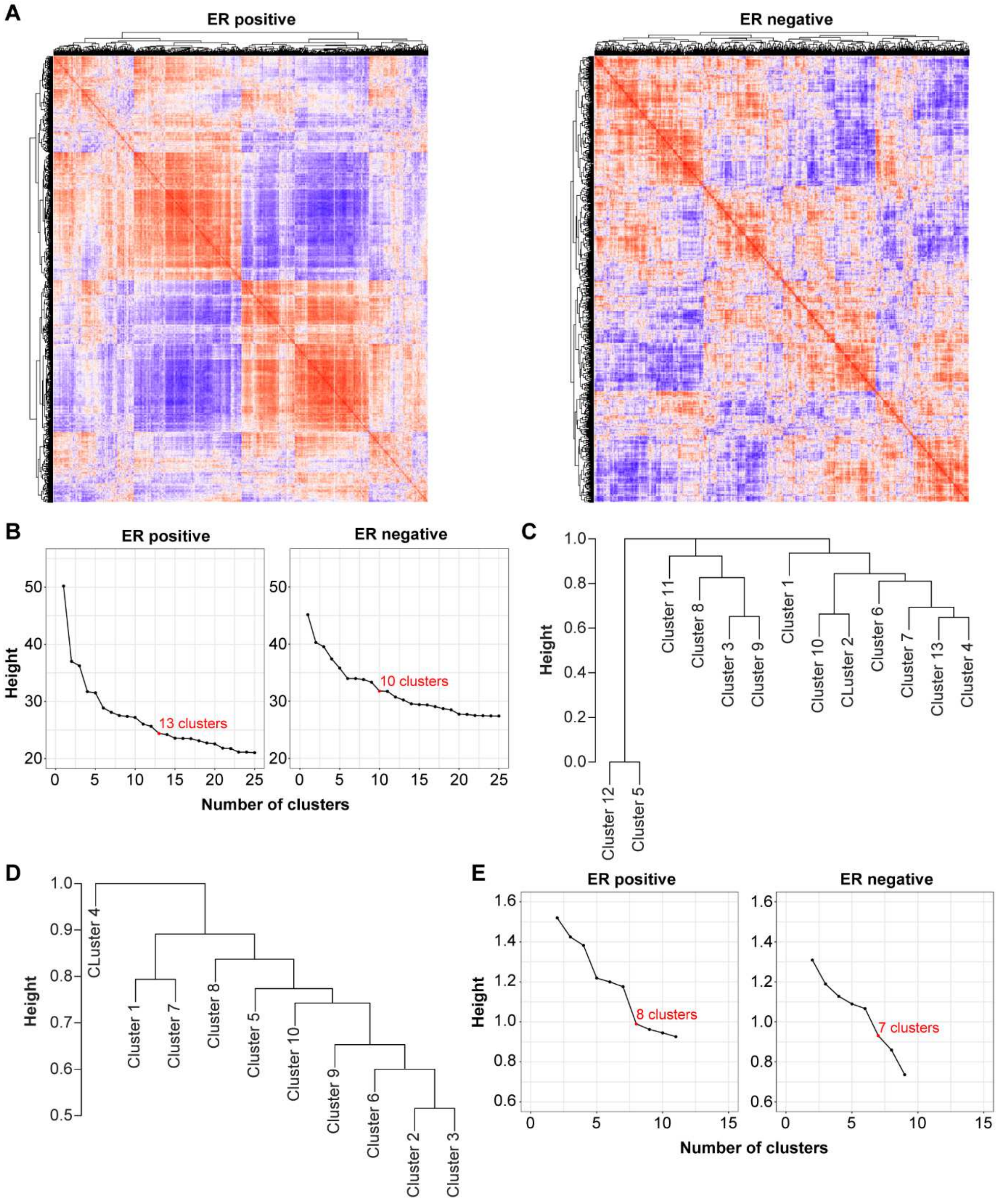

**Figure S12. Protein cluster regulation dependent on Estrogen Receptor status out of the DDA data layer.**

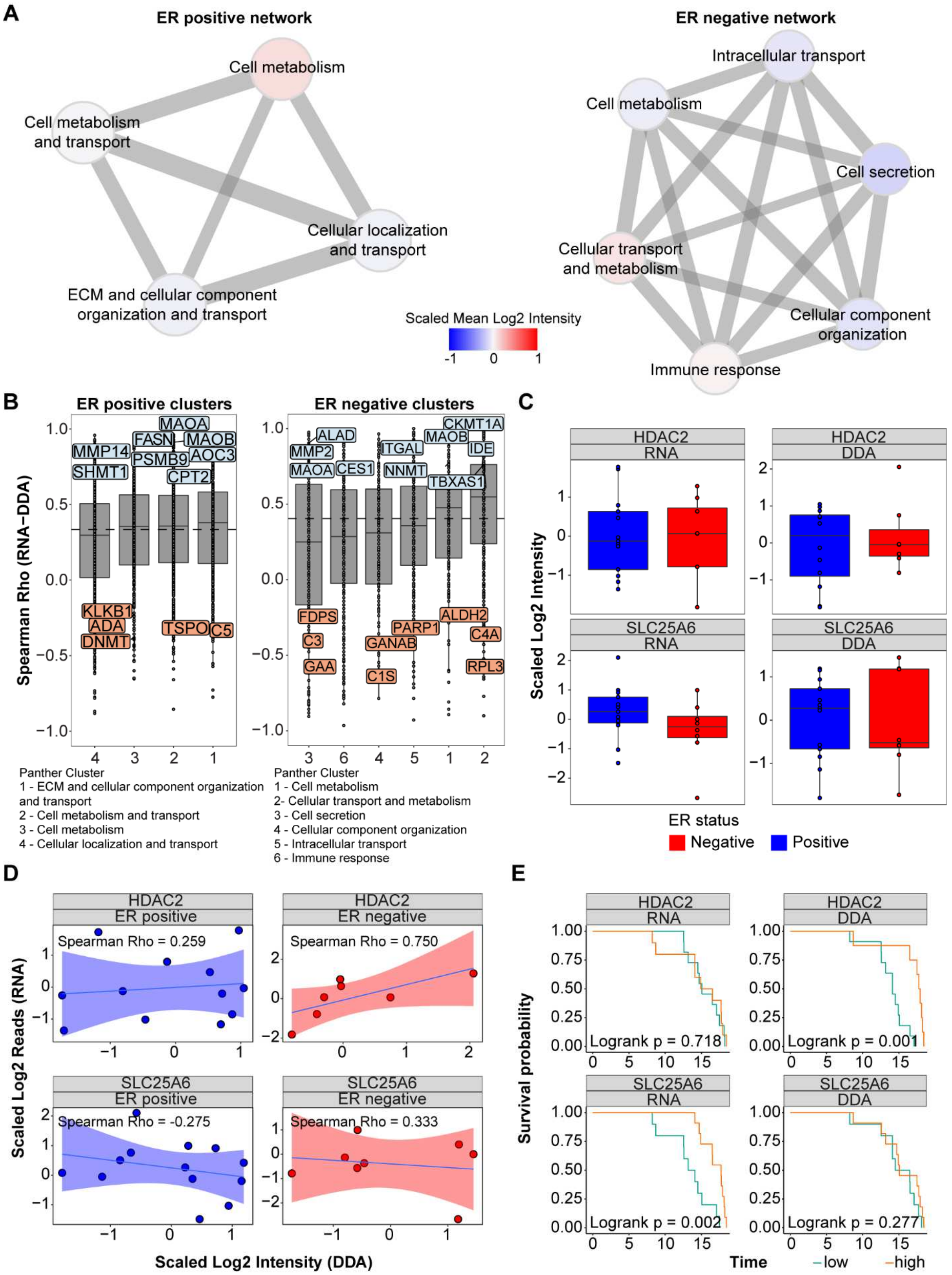
